# Decoding spine nanostructure in cultured neurons derived from mouse models of neuropsychiatric disorder reveals a schizophrenia-linked role for Ecrg4

**DOI:** 10.1101/2025.09.10.675343

**Authors:** Yutaro Kashiwagi, Qingrui Liu, Yasuhiro Go, Ryo Saito, Atsu Aiba, Takanobu Nakazawa, Shigeo Okabe

## Abstract

Dendritic spine dysfunction may contribute to the etiology and symptom expression of neuropsychiatric disorders. The intimate relationship between spine morphology and function suggests that decoding disease-related abnormalities from spine morphology can aid in developing synapse-targeted interventions. Here, we describe a population analysis of dendritic spine nanostructure applied to the objective grouping of multiple mouse models of neuropsychiatric disorders. This method has identified two major groups of spine phenotypes linked to schizophrenia and autism spectrum disorder (ASD). An increase in spine subpopulation with small volumes characterized the spines of schizophrenia-associated mouse models, whereas a spine subset with large volumes increased in ASD models. Schizophrenia-associated mouse models showed higher similarity in spine morphology, driven by reduced size and growth of nascent spines. The expression of Ecrg4, a gene encoding small secretory peptides, was increased in schizophrenia-associated mouse models, and functional studies confirmed its critical involvement in impaired spine dynamics and shape. These results suggest that population-level spine analysis provides rich insights into heterogeneous spine pathology, facilitating the identification of new molecular targets related to core synaptic dysfunction.

## Introduction

Information processing in the neocortex and hippocampus relies on fast excitatory synaptic transmission between pyramidal neurons. Many excitatory synapses that release glutamate project onto tiny structures protruding from dendritic shafts termed spines, and the morphological properties of these spines strongly influence synaptic transmission and function, including the activity-dependent synaptic plasticity implicated in learning and memory^1,2^. In general, larger spines contain a greater number of glutamate receptors and are functionally stronger^3^. The morphological parameters defining spine shape, such as length, radius, and volume, are also highly dynamic. For instance, small nascent spines can be enlarged by high-frequency stimuli, inducing long-term potentiation (LTP) of synaptic transmission^4,5^; alternatively, spine volume can be reduced by low-frequency stimuli that induce long-term depression (LTD)^5,6^. Due to these dynamic changes, spines on pyramidal neurons exhibit substantial morphological variation, even within the same dendritic branch. In addition to direct synaptic activation inducing LTP or LTD, spine morphology can be altered by cyclical hormonal changes, neuromodulators, and various activity-independent processes^7–10^. In turn, these changes in spine morphology are associated with marked changes in neural function and behavior.

Recent advances in super-resolution microscopy have provided new opportunities for the study of spine structural diversity and dynamics. An investigation using stimulated emission depletion (STED) imaging reported an increase in spine neck size after LTP, resulting in stronger electrical coupling between the spine head and the dendritic shaft^11^. A structured illumination microscopy (SIM) imaging study of cultured hippocampal neurons revealed expansion of the spine head convexity after LTP induction^12^, and suggested that this structural transformation may increase the adhesion of presynaptic and postsynaptic membranes. SIM imaging has also been applied to record the dynamic properties of spinules^13^, thin protrusions extending from existing spines that may function in secondary synapse formation. Collectively, these reports implicate spine nanostructural dynamics in multiple core synaptic properties and functions, including neurotransmission, mechanical stability, synaptogenesis, and plasticity.

Genetic mutations and variants that disrupt synaptic function are strongly implicated in the pathogenesis of neuropsychiatric disorders such as autism spectrum disorder (ASD) and schizophrenia^14–20^. Based on these genomic aberrations, numerous mouse models of ASD and schizophrenia have been established and subsequently shown to harbor abnormalities in both spine morphology and physiology^19,21–24^. Moreover, several studies have reported synaptic dysfunction in neurons differentiated from patient-derived pluripotent stem cells ^25–28^. Schizophrenia and ASD share several clinical features, suggesting common genetic risk factors and etiology^29,30^. However, a diametrical model emphasizes the distinct social-cognitive properties of these two major psychiatric disorders^31,32^. Both ASD and schizophrenia also show highly heterogeneous clinical phenotypes, potentially stemming from distinct pathogenic processes within disease categories. Therefore, in addition to clinical features, comparative biological phenotyping of multiple ASD and schizophrenia models may reveal shared as well as disease-specific pathogenic processes.

Genetic studies on ASD and schizophrenia have identified copy-number variants and protein-disrupting single-nucleotide variants that confer higher disease risk, and mouse models harboring these genetic abnormalities frequently exhibit behavioral and neurological features resembling those of the clinical conditions (termed endophenotypes)^19,33^. However, only a limited number of studies objectively compare phenotypes related to synaptic structure and function across disease models. An *in vivo* two-photon imaging study reported accelerated spine turnover in three genetically distinct mouse models of ASD^34^. Also, multiple mouse models of ASD were found to exhibit a similar imbalance in the activity of excitatory and inhibitory neurons^35^. These studies support the idea that distinct genetic mutations may induce similar synaptic and circuit phenotypes in ASD-associated mouse models. However, differences in synaptic properties between ASD and schizophrenia have not yet been comprehensively examined.

In this study, we developed an objective method for identifying population-level differences in spine nanostructure. This method, which has been applied to multiple mouse models of psychiatric disorders, identified two distinct groups corresponding to ASD and schizophrenia. An increase in a specific spine subpopulation with small volumes characterized the spines of mouse models of schizophrenia. In turn, ASD-related mouse models showed an opposite spine phenotype. The schizophrenia-related phenotype was associated with reduced size and growth of nascent spines and enhanced spine turnover. Further gene expression analysis identified the overexpression of Ecrg4, a gene encoding a precursor of hormone-like peptides, as a candidate mediator of this schizophrenia-associated spine phenotype. Population-level analysis of spine nanostructures is a powerful approach for understanding heterogeneous synaptic impairments in psychiatric disorders.

## Results

### Spine nanostructure imaging and generation of the spine density plot

We have designed a method for objectively comparing spine properties among multiple mouse models of psychiatric disorders at the population level using SIM imaging. Previous results demonstrated that high-resolution SIM images of dendritic spines in cultured hippocampal neurons can provide reliable information about spine size and morphological properties^12^. In the current study, the analytical procedures for SIM images were further improved for quantitative comparisons. Three-dimensional SIM images of dendritic spines were captured in DiI-stained cultured neurons derived from control and disease-model mouse hippocampus (Figure 1A). This staining procedure provided a better image quality than fluorescent protein-based labeling because of the higher fluorescence signal from the plasma membrane. SIM images were processed using custom-made scripts to identify and segment the individual dendritic spines (Figure 1B–D). We previously reported a set of scripts designed to isolate dendritic spines automatically^12^. However, the segmentation accuracy still necessitated manual inspection. In the newly developed scripts, we introduced a step to readjust the spine–shaft boundary, enabling fully automated measurement of multiple morphological parameters, including length, surface area, and volume. In addition, the isolated spines were further divided into 160-nm-thick longitudinal segments, yielding a total of 64 shape descriptors.

**Figure 1.**
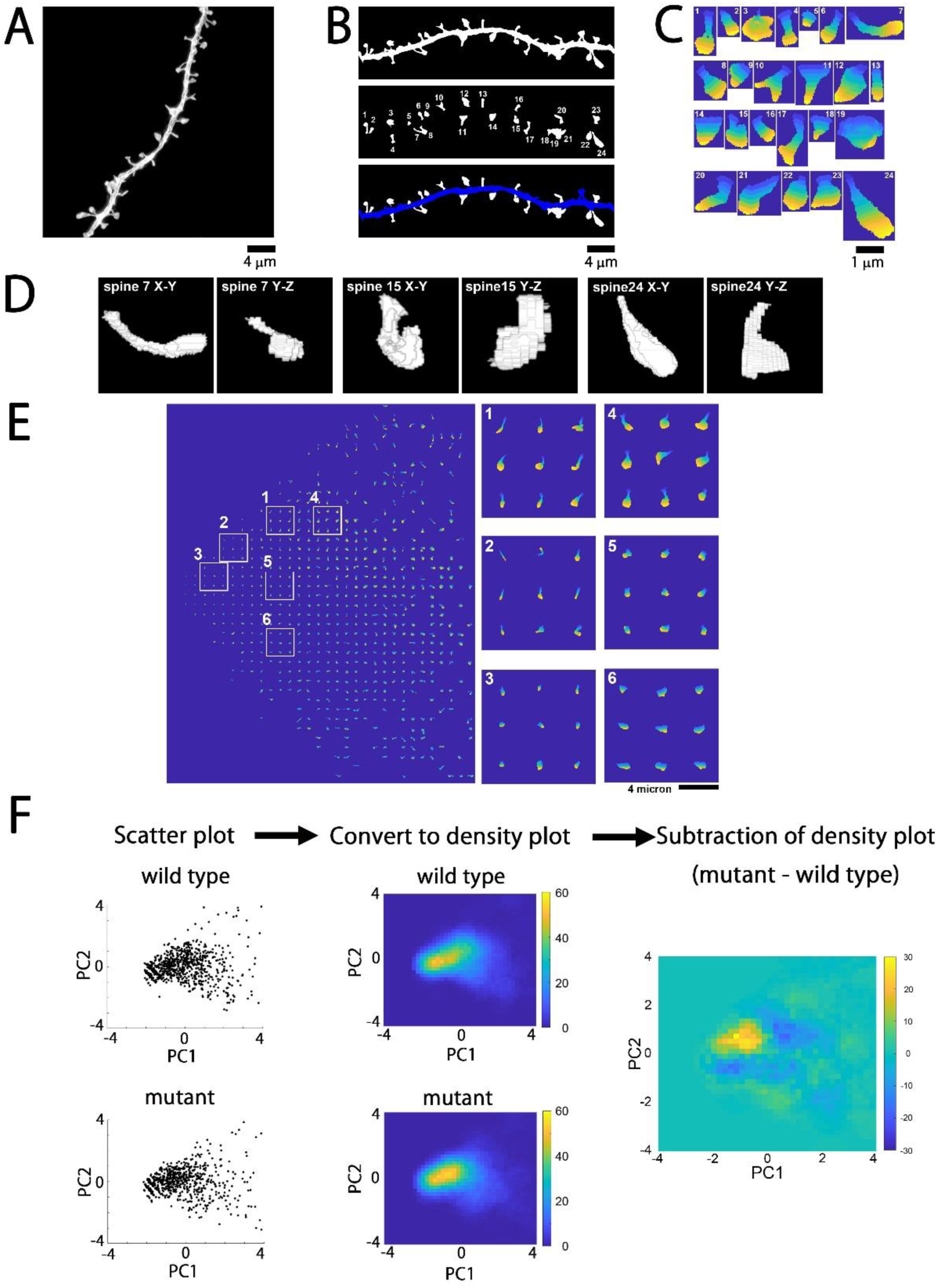
SIM imaging of dendritic spines, automated measurements of spine morphology, and generation of subtracted spine density plots for population-level analysis. (A) Original SIM image of a dendritic shaft stained with lipophilic membrane dye DiI. (B) Binarized image of the same dendrite (top) and segmented spines numbered from 1 to 24 (middle). The bottom image shows segmented spines (white) and the dendritic shaft (blue). (C) Enlarged images of the individual spines shown in (B). The pseudocolor images indicating the relative positions of the spine segments from the base (blue) to the tip (yellow). (D) 3D views of three spines (No. 7, 15, and 24) viewed from different angles. (E) Principal component analysis (PCA)-based dimensional reduction of spine characteristics plotted in the plane of principal components (PC)1 and PC2. (F) Process of generating a subtracted density plot. The scatter plot of spine distribution in the PC1–PC2 plane based on morphological parameters was converted into a density plot for each culture source (genotype or treatment group), and the corresponding plots were subtracted to reveal differences in spine morphology at the population level. Bars, 4 μm for A, B, and E, 1 μm for C.

Spine shape parameters measured by conventional methods are difficult to compare directly when images are obtained from multiple independent experiments. This problem is caused by several factors, but the main factors are variability in image intensity, signal-to-noise ratio, and image aberration introduced by unoptimized imaging conditions. To test the stability of SIM-based spine imaging, we compared the spine size distribution across four samples prepared at time points separated by >2 months (Supplementary Figure 1). The cumulative distribution curves of the four datasets showed extensive overlap, confirming the reproducibility of the SIM-based spine shape measurements using independent culture samples.

Another technical issue in comparing spine shape is the heterogeneity of spine shapes and the absence of defined subclasses. Population-level spine analysis, therefore, requires a new strategy that enables objective comparisons of large numbers of spines with a continuum of morphologic features. We previously demonstrated that principal component analysis (PCA) was effective for comparing high-dimensional spine shape features (descriptors) in a reduced-dimensional feature space^12^. Indeed, a PCA-based dimensional reduction of a large number of spines, based on their descriptors, generated a two-dimensional plot of spine shape that closely matched conventional spine subtypes, such as thin, mushroom, and stubby (Figure 1E). To objectively compare the two spine datasets (control and disease model), we first generated plots of relative spine densities (number per area in feature space) mapped to 80 × 80 square blocks (Figure 1F), and then subtracted the corresponding plot pairs defined by genotype (control vs. mutant) or culture conditions. The resulting subtracted density plots contain comprehensive information about the differences in relative spine numbers with similar shape properties. As expected, there was some variation in the density plot patterns across experiments. However, subtracting the distributional data from wild-type and mutant samples effectively canceled out these differences. Moreover, the difference between the two wild-type samples was generally smaller than that between wild-type and mutant samples (Supplementary Figure 2).

### Comparison and grouping of multiple mouse models based on morphological spine similarity

The subtracted density plots contain rich information about the population-level differences in spine shape between samples. Therefore, the generation and comparison of the plots for both ASD- and schizophrenia-associated mouse models will be useful for their unbiased grouping. We selected the following eight representative mouse models harboring either copy-number variation or single gene mutations associated with psychiatric disorders: male hemizygous or female homozygous mutant mice for neuroligin-3 R451C mutation (Nlgn3^R451C/(y^ ^or^ ^R451C)^)^36,37^, heterozygous Syngap1 mutant mice (Syngap1^+/−^)^38^, heterozygous POGZ mutant mice (POGZ^Q1038R/+^)^39^, mice paternal duplication of chromosome 7C (15q11-13^dup/+^) corresponding to human 15q11-13 duplication^40^, mice with heterozygous deletion of chromosome 16B2,3 (3q29^del/+^) corresponding to human 3q29 deletion^41^, mice heterozygous for deletion on chromosome 16qA13 (22q11.2^del/+^) corresponding to 1.5 Mb deletion on human 22q11.2^42^, mice heterozygous for Setd1a mutation (Setd1a^+/-^)^43^, and Ca^2+^/calmodulin-dependent protein kinase IIα kinase-dead mutant mice (CaMKIIα^K42R/K42R^)^44^. Previous studies on these mouse models confirmed the presence of multiple sensory, memory, and social endophenotypes of psychiatric disorders compared to controls. Of these, Nlgn3^R451C/(y^ ^or^ ^R451C)^, POGZ^Q1038R/+^, and 15q11-13^dup/+^ models mimic genetic variations found in ASD. Human mutations in the SYNGAP1 gene are associated with intellectual disability and autistic behaviors. The 3q29^del/+^, 22q11.2^del/+^, and Setd1a^+/-^ models harbor genetic mutations associated with higher schizophrenia risk. CaMKII-related signaling pathway disruption has been implicated in the working memory deficits found in schizophrenia patients^45,46^. CAMK2A mutations in humans are linked to multiple mental disorders, including developmental disorders, ASD, and schizophrenia^47^. The K42R mutation of CAMK2A does not correspond to any known human genetic variant, but the CAMK2A R8H mutation is linked to schizophrenia^48^. Both R8H and K42R mutations in the N-terminal domain of CaMKIIα eliminate kinase activity; these mutations may have a similar impact on human mental disorders. Taken together, these mouse models cover a broad spectrum of human genetic aberrations associated with psychiatric disorders. Direct comparison of spine phenotypes could therefore provide important clues to underlying shared and disease-specific synaptic pathologies.

For these comparisons, we prepared three independent cultures of embryonic hippocampi from each control–mutant pair and obtained 700–1,500 3D-SIM spine images 3 weeks after plating for both mutant and control samples (Nlgn3^R451C/(y^ ^or^ ^R451C)^, n = 1,134 and 1,204; Syngap1^+/−^, n = 991 and 1,371; POGZ^Q1038R/+^, n = 1,341 and 1,271; 15q11-13^dup/+^, n = 1,208 and 1,099; 3q29^del/+^, n = 1,429 and 1,408; 22q11.2^del/+^, n = 1,143 and 914; Setd1a^+/-^, n = 1,381 and 1,405; CaMKIIα^K42R/K42R^; n = 880 and 763, control and mutant, respectively, Supplementary Table 1). Initial comparison of the cumulative frequency plots for spine length, spine surface area, and spine volume alone did not provide sufficient information to infer their structural similarity (Supplementary Figure 3), indicating that simple structural parameters are not effective in identifying disease-specific spine shape characteristics. We next generated subtracted density plots from 8 mouse models. These plots were variable, but there were certain similarities between the specific mutants (Figure 2A). We evaluated pairwise similarities by calculating the 2D cross-correlations of the subtracted density plots. The matrix of these 2D cross-correlation values yielded two major groups with high within-group similarity (Figure 2B), and this grouping was further confirmed by unbiased clustering analysis (Figure 2C). The ASD models Nlgn3^R451C/(y^ ^or^ ^R451C)^, Syngap1^+/−,^ and POGZ^Q1038R/+^ were in the first group, while the second group contained the schizophrenia models 3q29^del/+^, 22q11.2^del/+^, and Setd1a^+/-^. This result is striking, as it indicates that information about spine shape alone can distinguish whether the culture preparation is derived from ASD- or schizophrenia-associated mouse models. It is also of note that there were greater similarities among the schizophrenia-associated mouse models than among the ASD-associated mouse models. This finding suggests convergence in spine pathophysiology across schizophrenia-associated mouse models.

**Figure 2.**
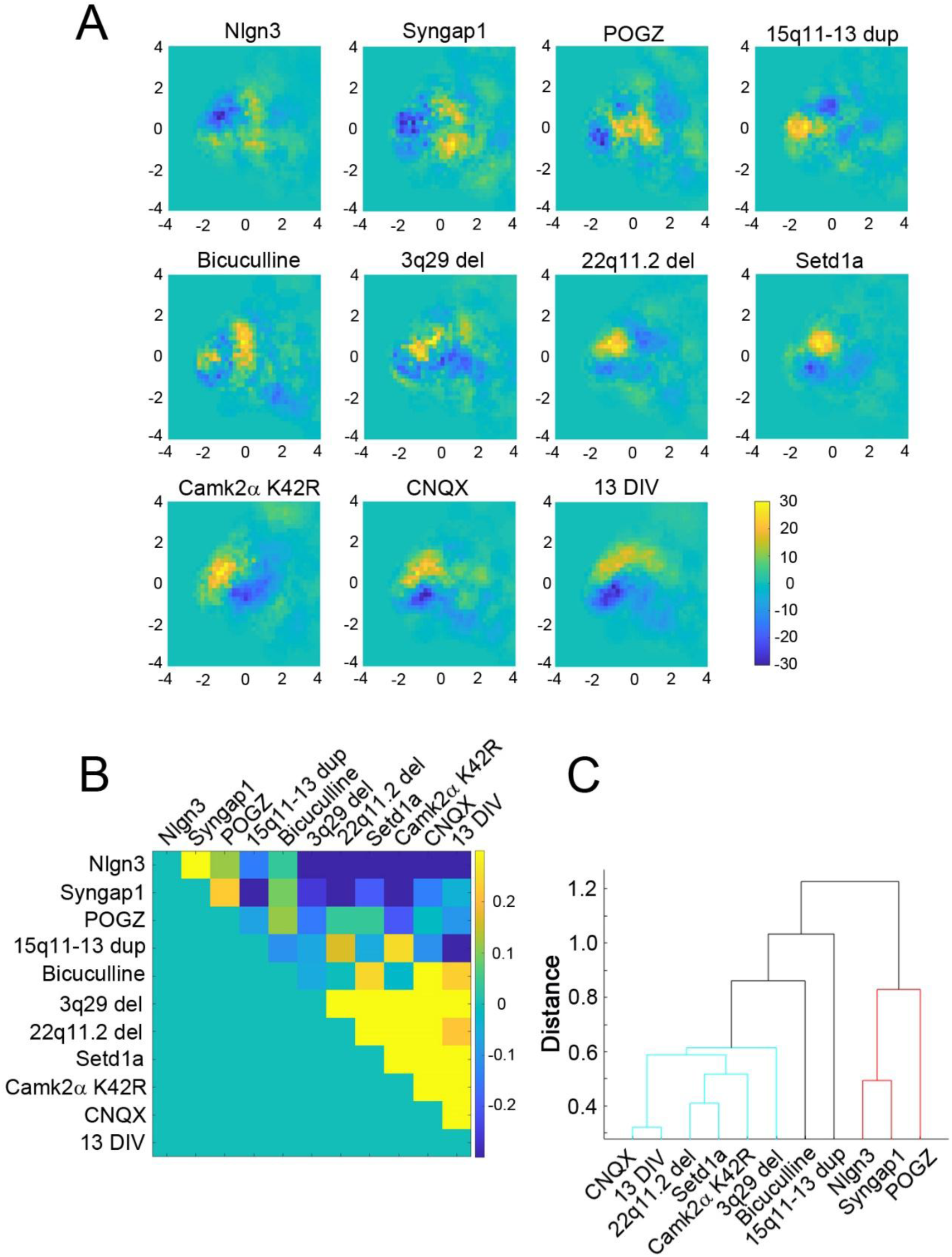
Spine population profiles for each model and corresponding control mouse line presented as subtracted density plots. (A) The subtracted density plots for eight disease mouse models (Nlgn3^R451C/(y^ ^or^ ^R451C)^, Syngap1^+/−^, POGZ^Q1038R/+^, 15q11-13^dup/+^, 3q29^del/+^, 22q11.2^del/+^, Setd1a^+/-^, and CaMKIIα^K42R/K42R^) and three different culture conditions (immature culture at 13 DIV, AMPA glutamate receptor blocker CNQX treatment, and GABA_A_ receptor blocker bicuculine treatment). The areas with a higher density of spines from mutant disease model mice are shown in yellow, and the areas with reduced density are shown in blue. The total number of spines (first number, n) analyzed from control and mutant mouse neurons and the corresponding number of dendrites (number in parentheses) are as follows: Nlgn3^R451C/(y^ ^or^ ^R451C)^; n = 1,134 (58) and 1,204 (59), Syngap1^+/−^; n = 991 (65) and 1,371 (83), POGZ^Q1038R/+^; n = 1,341 (72) and 1,271 (72), 15q11-13^dup/+^; n = 1, 208 (68) and 1,099 (63), 3q29^del/+^; n = 1,429 (66) and 1,408 (66), 22q11.2^del/+^; n = 1,143 (66) and 914 (63), Setd1a^+/-^; n = 1,381 (71) and 1,405 (71), CaMKIIα^K42R/K42R^; n = 880 (58) and 763 (54). All data are from three independent culture preparations. (B) Matrix of the 2D cross-correlations among subtracted density plots. In the lower right area, a spine group showing similar morphological changes can be identified. This group corresponded to the mouse models of schizophrenia. (C) Unbiased clustering of spine samples showing two distinct groups corresponding to schizophrenia (cyan) and ASD (red).

In addition to the eight mouse models of psychiatric disorders, we generated subtracted density plots for three additional conditions: immature culture samples (13 days in vitro, DIV), treatment with the AMPA subtype glutamate receptor blocker CNQX, and treatment with the GABA_A_ receptor blocker bicuculline (Figure 2A). The subtracted density plots from the 13 DIV culture and the CNQX treatment were similar to those of schizophrenia-associated mouse models. This finding suggests that the hippocampal synapses in mouse models of schizophrenia are immature and inactive. In contrast, the subtracted density plot of the bicuculline-treated sample was distinct from that of both the schizophrenia and ASD-associated mouse models.

### Distinct spine shapes between ASD- and schizophrenia-associated mouse models

The subtracted density plots provide information on the structural properties of specific spine subsets that either increased or decreased in disease mouse models. We first defined the four areas within the PC1-PC2 plane with distinct shapes: (1) small and short, (2) small and long, (3) large and short, and (4) large and long (Figure 3A and B). We also identified the wildtype-enriched area (blue areas in Figure 3C and D) and the mutant-enriched area (yellow areas in Figure 3C and D) in the same projection plane. The comparison of these two types of areas suggests that larger spines are enriched in the Nlgn3^R451C/(y^ ^or^ ^R451C)^ mouse model, while small spines are enriched in the 22q11.2^del/+^ mouse model. Next, we calculated the normalized spine counts in the four areas for both wild-type and mutant samples and obtained the relative abundance (mutant/wild-type) for each area (Figure 3E and F). The average of three independent culture preparations confirmed the opposite trend between Nlgn3^R451C/(y^ ^or^ ^R451C)^ and 22q11.2^del/+^.

**Figure 3.**
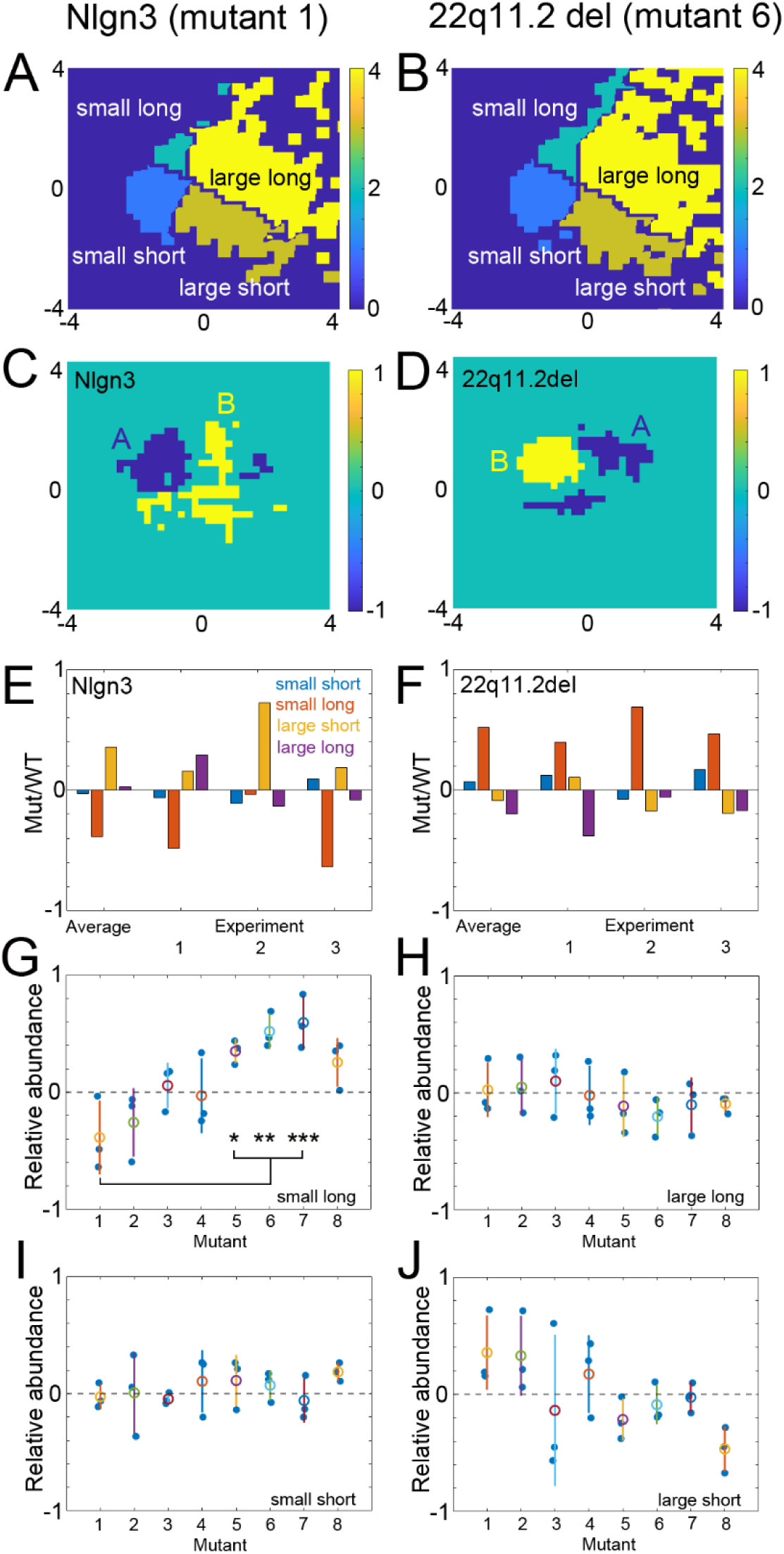
Distinct morphological properties of spines in cultured neurons derived from schizophrenia model mice compared with ASD model mice. (A, C, and E) Relative enrichment of four different subgroups of spines in Nlgn3^R451C/(y^ ^or^ ^R451C)^ neurons compared with wild-type neurons. In the projection plane of PC1 and PC2, four areas with distinct structural properties, namely (1) small and short, (2) small and long, (3) large and short, and (4) large and long, were defined (A). The map of areas with large differences (> 3 × SD) between control and mutant mice in spine number per area in the feature space (C) indicates that the area enriched with wild-type spines (area B, yellow region) overlaps with the area of large spines. The area enriched with the mutant spines overlaps with the area of small spines (area A, blue region). The relative enrichment of mutant spines was summarized in (E). (B, D, and F) Similar analysis of mutant spine enrichment in 22q11.2^del/+^ neurons. The map of spine distribution with four different properties (B), the map of areas enriched with wild-type or mutant spines (D), and the graph of relative enrichment of mutant spines (F) show that the mutant neurons were enriched with small and long spines. (G-J) Relative abundance of the four subgroups of spines in 8 mutant mouse models (1: Nlgn3^R451C/(y^ ^or^ ^R451C)^, 2: Syngap1^+/−^, 3: POGZ^Q1038R/+^, 4: 15q11-13^dup/+^, 5: 3q29^del/+^, 6: 22q11.2^del/+^, 7: Setd1a^+/-^, 8: CaMKIIα^K42R/K42R^). Three independent culture experiments with paired wild-type and mutant samples were performed. Group comparisons were performed using one-way ANOVA followed by Tukey’s post hoc test. (* p < 0.05, ** p < 0.01, *** p < 0.005).

Figure 3G-J summarizes the relative abundance of mutant spines in four areas with distinct spine shapes. We found that the relative abundance of mutant spines in area 2 (small and long spines) differed significantly between genotypes (ANOVA, p < 0.001). Group comparisons by Tukey’s post hoc test indicate significant differences between Nlgn3^R451C/(y^ ^or^ ^R451C)^ mutant and three schizophrenia-related mutants; 3q29^del/+^, 22q11.2^del/+^, and Setd1a^+/-^ (Figure 3G). The opposite trend between areas 2 (small and long spines) and 3 (large and short spines) further confirms distinct spine-shape properties in ASD- and schizophrenia-related mouse models. The areas within the subtracted density plots with enriched control or mutant spines illustrate the characteristics of spine populations across mouse models (Supplementary Figures 4 and 5). The shape profiles of spines confirmed smaller spine volume of schizophrenia-related mouse models (Supplementary Figure 6).

### Altered initial growth of dendritic spines in mouse models of schizophrenia

The subtracted density plots reveal common changes in the spine population across schizophrenia-associated mouse models. The mutant-spine-enriched areas also contain small, elongated spines, suggesting dysfunction in either the generation of nascent spines or the shrinkage of existing spines. To examine potential abnormalities in spine dynamics, we performed time-lapse imaging of hippocampal neuron cultures derived from two schizophrenia-associated mouse models, 22q11.2^del/+^ and Setd1a^+/-^, and the Nlgn3^R451C/(y^ ^or^ ^R451C)^ mouse model as a reference (Figure 4).

**Figure 4.**
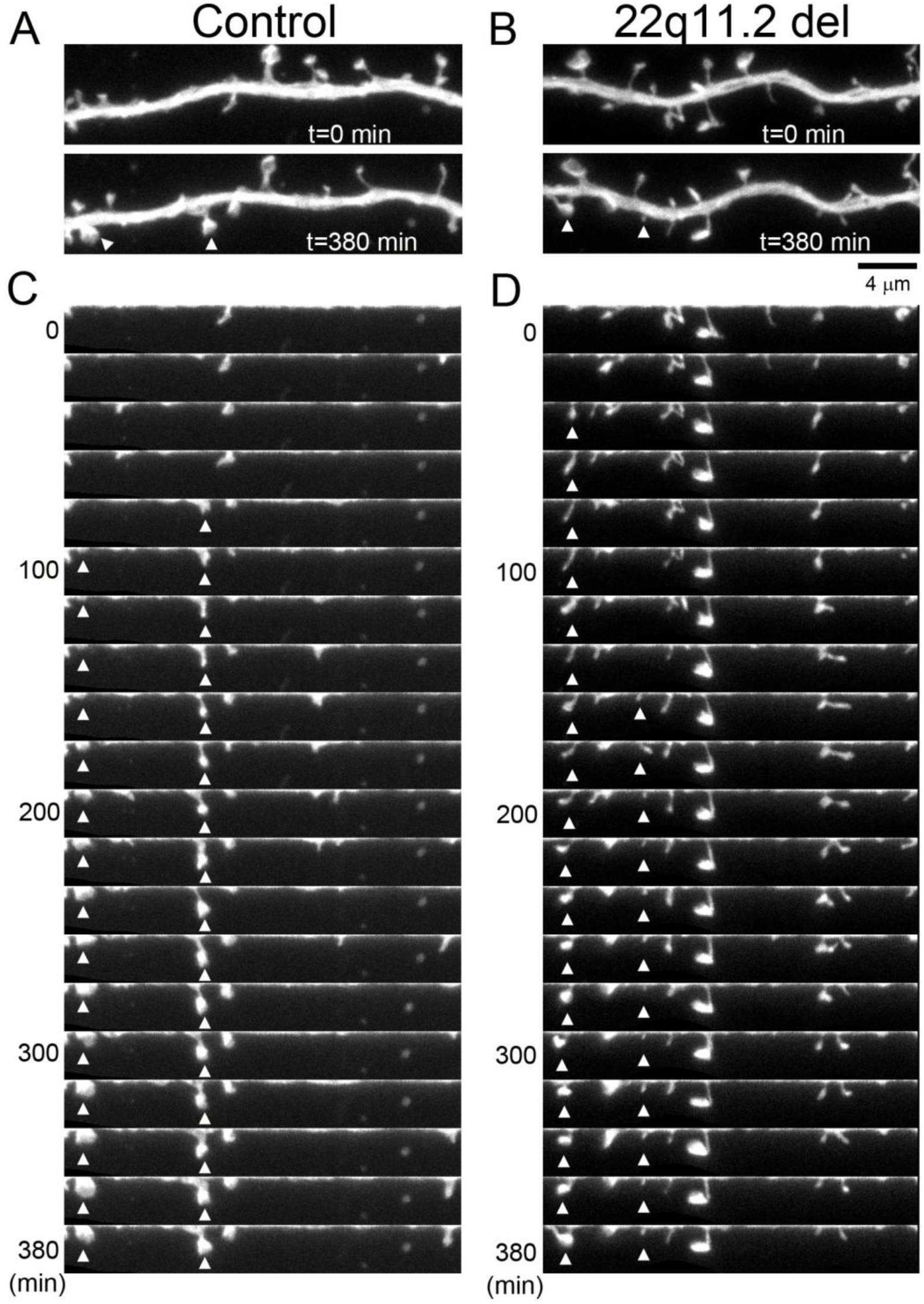
Time-lapse imaging of neurons derived from 22q11.2^del/+^ mice and corresponding control mice. (A and B) Images of dendritic segments from control neurons (A) and 22q11.2^del/+^ neurons (B) at two different time points. (C and D) Montages of time-lapse images from control neurons (C) and 22q11.2^del/+^ neurons (D). The curved dendrites were straightened, revealing newly formed spines (arrowheads) as fluorescent objects appearing at the edge of the dendritic shafts. Bar, 4 μm.

A linear mixed-effects model indicated significant effects of genotype in the increase of spine turnover rate in both 22q11.2^del/+^ and Setd1a^+/-^ models (F(1,25) = 5.79, p = 0.024 for 22q11.2^del/+^ and F(1,22) = 7.33, p = 0.013 for Setd1a^+/-^). In contrast, no significant effects were observed in the Nlgn3^R451C/(y^ ^or^ ^R451C)^ model (Figure 5A-C). There was a significant effect of genotype in both 22q11.2^del/+^ and Setd1a^+/-^ models also for the prolonged lifetime of transient spines (a linear mixed-effects model: 22q11.2^del/+^; F(1,336)=5.33, p=0.022, Setd1a^+/-^; F(1,282)=6.38, p=0.012), with no significant difference in Nlgn3^R451C/(y^ ^or^ ^R451C)^ model (Figure 5D-F).

**Figure 5.**
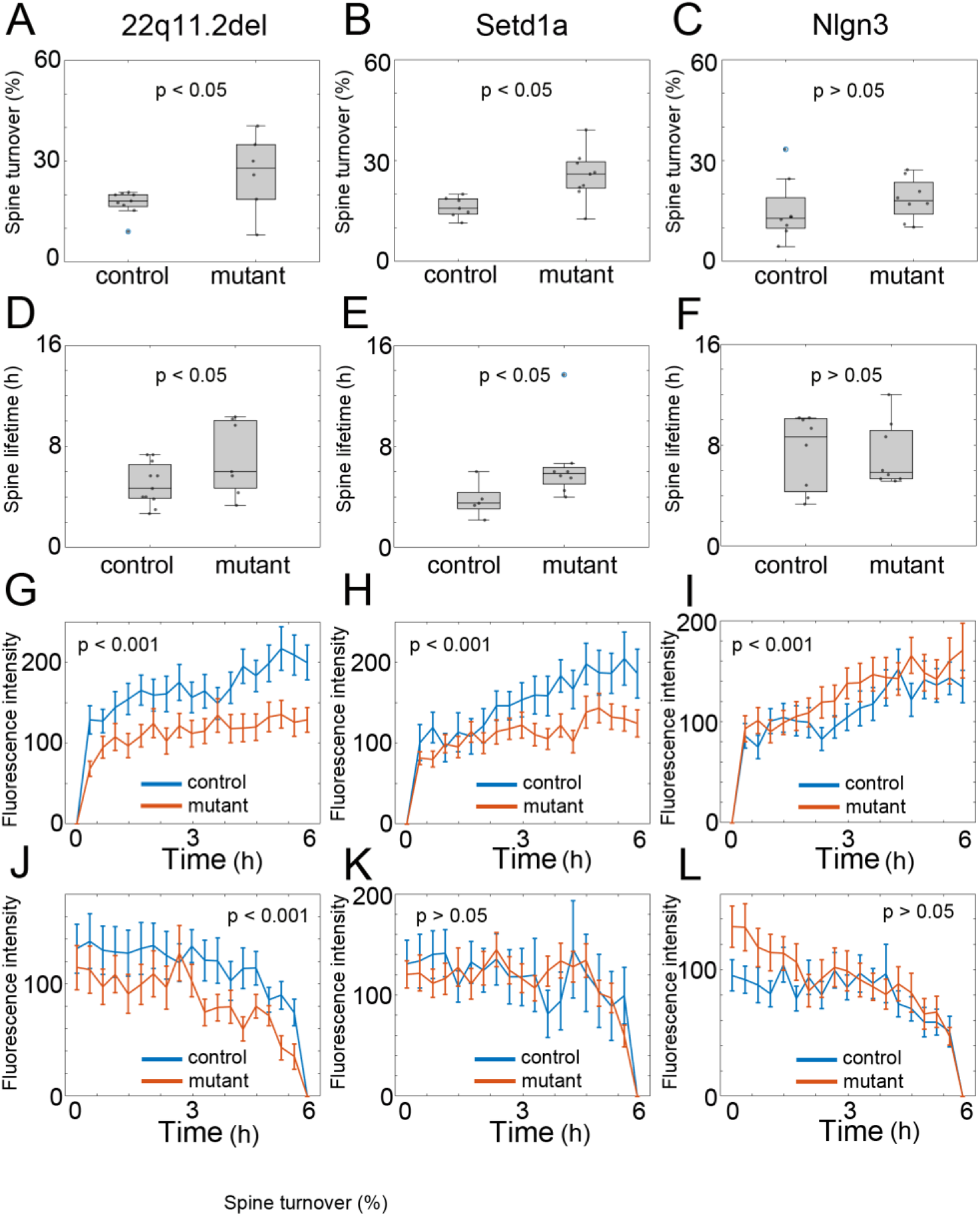
Turnover rate, lifetime, and growth/shrinkage profiles of dendritic spines in cultured neurons derived from 22q11.2^del/+^, Setd1a^+/-^, and Nlgn3^R451C/(y^ ^or^ ^R451C)^ mice, as well as the corresponding controls. (A-C) Spine turnover rates in the three mutant mouse models compared to the controls. Spine turnover rates were analyzed using a linear mixed-effects model with genotype as a fixed effect and plate, cell, and dendrite as nested random effects, showing a significant effect of genotype only in 22q11.2^del/+^ and Setd1a^+/-^ mice (n = 16 dendrites/9 cells/5 plates in control and 11 dendrites/6 cells/4 plates in 22q11.2^del/+^, n = 10 dendrites/7 cells/4 plates in control and n = 14 dendrites/9 cells/6 plates in Setd1a^+/-^, n = 8 dendrites/8 cells/3 plates in control and n = 8 dendrites/8 cells/3 plates in Nlgn3^R451C/(y^ ^or^ ^R451C)^). (D-F) Lifetimes of transient spines for the three mutant mouse models compared with corresponding controls. Spine lifetime was analyzed using a linear mixed-effects model accounting for the hierarchical structure of the data (spines nested within dendrites, cells, and culture plates). The analysis revealed a significant effect of genotype only in 22q11.2^del/+^ and Setd1a^+/-^neurons (n = 186 spines/11 dendrites/7 cells/5 plates in control and n = 166 spines/7 dendrites/4 cells/4 plates in 22q11.2^del/+^, n = 82 spines/5 dendrites/5 cells/4 plates in control and n = 202 spines/8 dendrites/8 cells/5 plates in Setd1a^+/-^, n = 98 spines/8 dendrites/8 cells/3 plates in control and n = 125 spines/8 dendrites/8 cells/3 platess in Nlgn3^R451C/(y^ ^or^ ^R451C)^). (G-I) Temporal patterns of spine growth (n = 60 spines/11 neurons/7 cells/5 plates in control and n = 50 spines/7 dendrites/4 cells/4 plates in 22q11.2^del/+^, n = 25 spines/5 dendrites/5 cells/4 plates in control and n = 65 spines /8 dendrites/8 cells/5 plates in Setd1a^+/-^, n = 43 spines/8 dendrites/8 cells/3 plates in control and n = 41 spines/7 dendrites/7 cells/2 plates in Nlgn3^R451C/(y^ ^or^ ^R451C)^). Spine volume trajectories were analyzed using linear mixed-effects models incorporating nested random effects (spine within dendrite within cell within culture plate) to account for the hierarchical structure of the data. Newly formed spines in both 22q11.2^del/+^ and Setd1a^+/-^ neurons were significantly smaller than those in wild-type neurons. In contrast, newly formed spines in Nlgn3^R451C/(y^ ^or^ ^R451C)^ neurons were significantly larger than those in wild-type neurons. (J-L) Temporal patterns of spine shrinkage (n = 39 spines/10 dendrites/6 cells/5 plates in control and n = 37 spines/7 dendrites/4 cells/4 plates in 22q11.2^del/+^, n = 15 spines/5 dendrites/5 cells/4 plates in control and n = 55 spines/8 dendrites/8 cells/5 plates in Setd1a^+/-^, n = 28 spines/8 dendrites/8 cells/3 plates in control and n = 35 spines/8 dendrites/8 cells/3 plates Nlgn3^R451C/(y^ ^or^ ^R451C)^ neurons). Spine volume trajectories were analyzed using linear mixed-effects models as in (G-I). In the 22q11.2^del/+^ neurons, spines undergoing elimination were significantly smaller than those in wild-type neurons. This effect was not detected in Setd1a^+/-^ and Nlgn3^R451C/(y or R451C)^ neurons.

We further compared the initial increase in spine volume between genotypes (Figure 5G-I). Linear mixed-effects models incorporating nested random effects revealed significantly smaller initial spine volumes in both 22q11.2^del/+^ and Setd1a^+/-^models (genotype effect: p < 0.001 for 22q11.2^del/+^ and p < 10⁻⁷ for Setd1a^+/-^). The spines in both mutants also displayed a significant reduction in spine volume increase (p < 0.001). In contrast, newly formed spines in the Nlgn3^R451C/(y^ ^or^ ^R451C)^ neurons were significantly larger than those in wild-type neurons (p < 10⁻⁴) with preserved time-course of spine growth.

Similar analyses of spine volume reduction before spine loss using linear mixed-effects models identified significantly smaller initial spine size in the 22q11.2^del/+^ model (p < 10⁻^6^), while no significant differences in the initial spine volume were found for the Setd1a^+/-^ model (Figure 5J-L). The temporal trajectories of spine shrinkage before their loss were not significantly altered in both 22q11.2^del/+^ and Setd1a^+/-^ models. The Nlgn3^R451C/(y^ ^or^ ^R451C)^ model showed a significantly different time-course of spine shrinkage (p < 0.05), while the initial spine size was not altered. Taken together, these results suggest that the smaller size of the initial spine, slower growth of nascent spines, and enhanced spine turnover contribute to the development of schizophrenia-related spine phenotypes.

To further examine the relationship between the spine growth rate and lifetime, we performed simulations using the computational model shown in Figure 6A. In this model, new spines appear randomly along the dendritic shaft according to a density-dependent probability function, then increase in volume with speed V1 (phase 1), are eliminated with probability P1 (phase 2), and are converted to stable spines upon reaching a volume threshold (phase 3). These stable spines are subsequently destabilized with low probability P2 (phase 4) and start to shrink with speed V2 (phase 5). When these shrinking spines reach a threshold, they are permanently removed. We did not incorporate the initial spine size as a model parameter because of its large variability, which made parameter optimization difficult. This model was run over 10 days of neuronal differentiation in culture (corresponding to the period of spine density increase from 8 to 17 DIV) with 625 different combinations of V1, V2, P1, and P2 to identify the combination with the best fit to the experimental data for spine turnover of 15–18 DIV neurons derived from mutants 22q11.2^del/+^, Setd1a^+/-^, and Nlgn3^R451C/(y^ ^or^ ^R451C)^ over 24 h (Figure 6B and Supplementary Figure 7A). The best-fitted models all yielded spine lifespan distributions matching observations of the three mouse mutants (Figure 6C). The two characteristics of the spines in 22q11.2^del/+^ and Setd1a^+/-^ were also preserved in the best-fitted model. First, spine density in the schizophrenia-associated mouse models was lower in both the simulation and experimental data (simulation: 0.57 ± 0.029 /μm and 0.40 ± 0.027 /μm for the control and the best-fitted for schizophrenia models, shown in Supplementary Figure 7B, experiment; 0.69 ± 0.17 /μm and 0.56 ± 0.20 /μm for the control and 22q11.2^del/+^, 0.70 ± 0.15 /μm and 0.64 ± 0.16 /μm for the control and Setd1a^+/-^). Second, the size of transient spines was smaller in schizophrenia-associated mouse models in the simulation (0.33 ± 0.033 and 0.20 ± 0.024 (a.u.) for the control and the best-fitted schizophrenia, Supplementary Figure 7C), consistent with data on spine nanostructure (Figure 3G and J).

**Figure 6.**
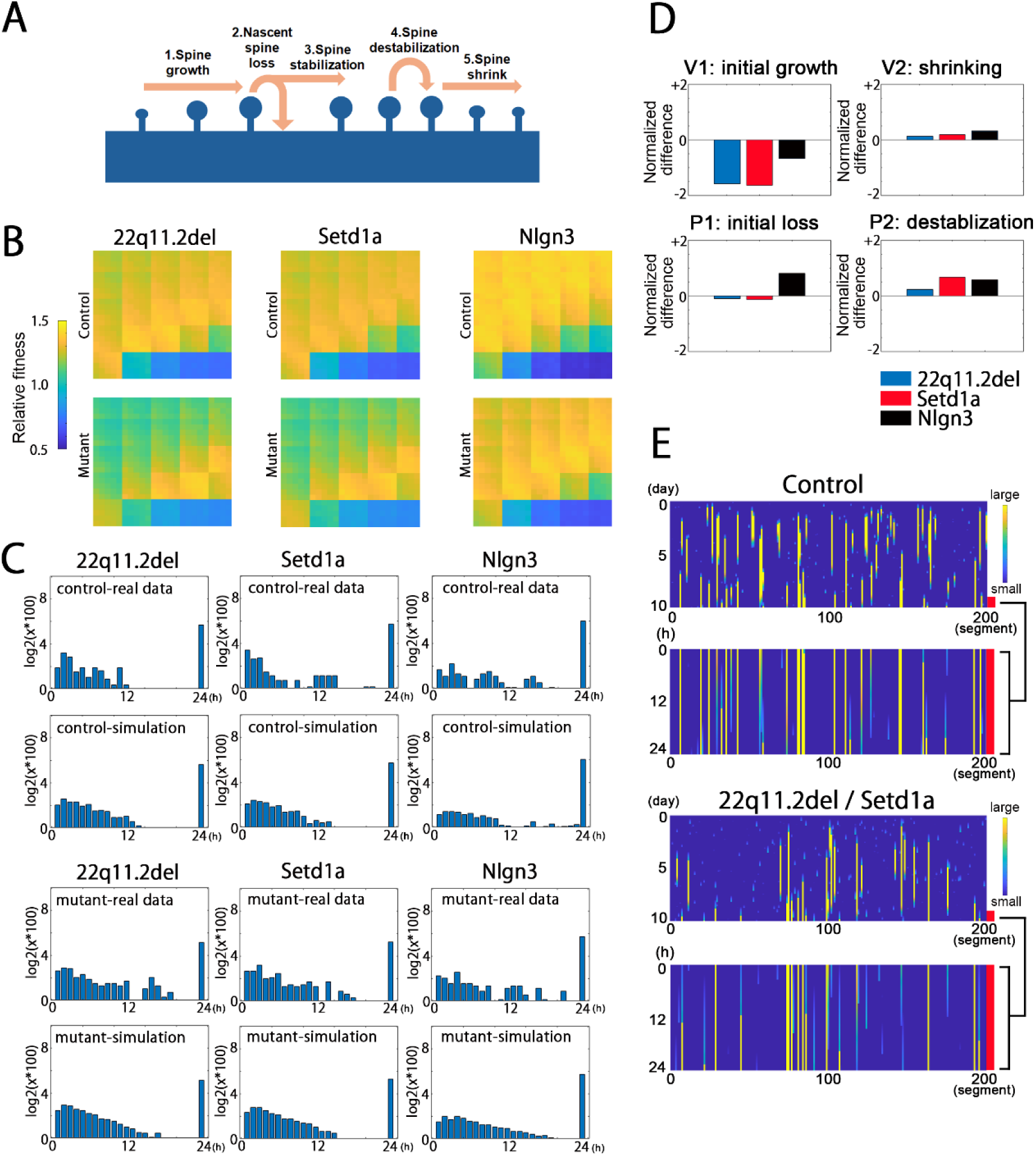
Simulations of long-term spine turnover. (A) The model of spine dynamics. Five successive phases of spine state transitions were defined. Phase 1: newly formed spines grow at speed V1. Phase 2: nascent spines are eliminated with probability P1. Phase 3: nascent spines are stabilized when volume reaches an upper threshold. Phase 4: stable spines are destabilized with probability P2. Phase 5: spines shrinking at a rate V2 are lost after reaching a lower threshold. (B) Pseudocolor maps of 625 different combinations of these parameters to identify those best fitting the experimental data. The color-coded values indicate the fitness of the simulation results to both the spine lifetime distribution profile and the turnover rate. (C) Frequency histograms of spine lifetimes for the three mouse models and controls (experimental data), and the results of simulations. The bin with a lifetime of 24 h corresponds to the spines that persisted throughout the imaging period. (D) Differences in V1, V2, P1, and P2 between mutant mice and controls, expressed as differences in the four parameters normalized by the control condition values. (E) Plots of individual spine turnover simulated using parameters that best fit the experimental data from control and schizophrenia-associated mouse models. The upper plots show the progression of spine formation along 200 dendritic segments (50 μm) over 10 days. The lower plots show the enlargement of the last 24 h of spine turnover. Color indicates spine size.

After the selection of the best-fitted parameters for three mouse models, we compared the optimal values of V1, V2, P1, and P2 between mouse models (Figure 6D). The nascent spine volume increase (V1) was slower in schizophrenia models than in corresponding controls. In turn, the probability of nascent spine loss (P1) was higher in the ASD model. In all three disease models, the probability of spine destabilization (P2) was moderately higher than that in the corresponding controls. Additional simulation results with different P2 values indicate that P2 has a small effect on the overall pattern of spine dynamics. These results suggest that a slow spine volume increase in schizophrenia model mice results in the accumulation of nascent spines with smaller volumes. On the other hand, the spines of ASD model mice grow at a rate similar to that of control spines, but are more likely to be eliminated both in the nascent stage and after maturation.

Simulated trajectory plots of individual spines along the dendritic shaft, using the best-fit parameters, also reflected the differential patterns of spine turnover in culture preparations derived from control and schizophrenia model mice (Figure 6E). In culture preparations derived from schizophrenia-associated mouse models, there were fewer spines with a lifetime longer than 2 or 3 days, while there was a greater number of spines formed and eliminated within 24 h. This increase in the number of less stable spines explains the pattern of spine lifetimes shown in Figure 5D-E. In summary, these simulations suggest that a reduced spine maturation rate can account for three key properties of schizophrenia-related spines: the greater number of small-volume spines, enhanced turnover, and reduced spine density.

### Screening of molecules responsible for altered spine properties in schizophrenia-associated mouse models

Both SIM-based imaging and time-lapse imaging suggest that the initial growth phase of spines is slower in schizophrenia model mice. To identify candidate genes that may contribute to this growth impairment, we performed RNA sequencing of libraries derived from the cultured neurons of eight psychiatric disorder mouse models and corresponding controls, and filtered the results for differentially expressed genes (DEGs) (Supplementary Table 2 and 3). The DEGs present in three or more schizophrenia-related mouse models are listed in Supplementary Figure 8. In the hippocampi of schizophrenia model mice, the expression levels of Met, Osr1, Arhgap15, and Rnf170-ps were downregulated, whereas those of Cip4, Ecrg4, and Npas4 were upregulated. The majority of these DEGs have been previously implicated in the pathology of schizophrenia (Met)^49^, synaptic plasticity (Npas4)^50,51^, or signaling pathways in the nervous system (Arhgap15^52^, Cip4^53^, and Ecrg4^54^).

If these genes do regulate spines in schizophrenia-associated mouse models, manipulation of these genes in wild-type neurons may induce a spine phenotype similar to that observed in these models. To test this possibility, we downregulated Met and Arhgap15, and upregulated Cip4, Ecrg4, and Npas4 in separate wild-type hippocampal neuron cultures and measured spine dynamics using time-lapse imaging (Figure 7A and B, Supplementary Figure 9). The application of a linear mixed-effects model to knockdown experiments revealed no significant effect of treatment (F(2,6) = 0.29, p = 0.76). In the overexpression experiment, the analysis using a linear mixed-effects model showed a significant main effect of treatment (F(3,8) = 4.59, p = 0.038), with a significant increase in spine turnover rate by Ecrg4 overexpression (β = 0.49 ± 0.16 SE, t(8) = 3.16, p = 0.013). Thus, the aberrant expression of Ecrg4 may be related to the spine phenotype of schizophrenia model mice.

**Figure 7.**
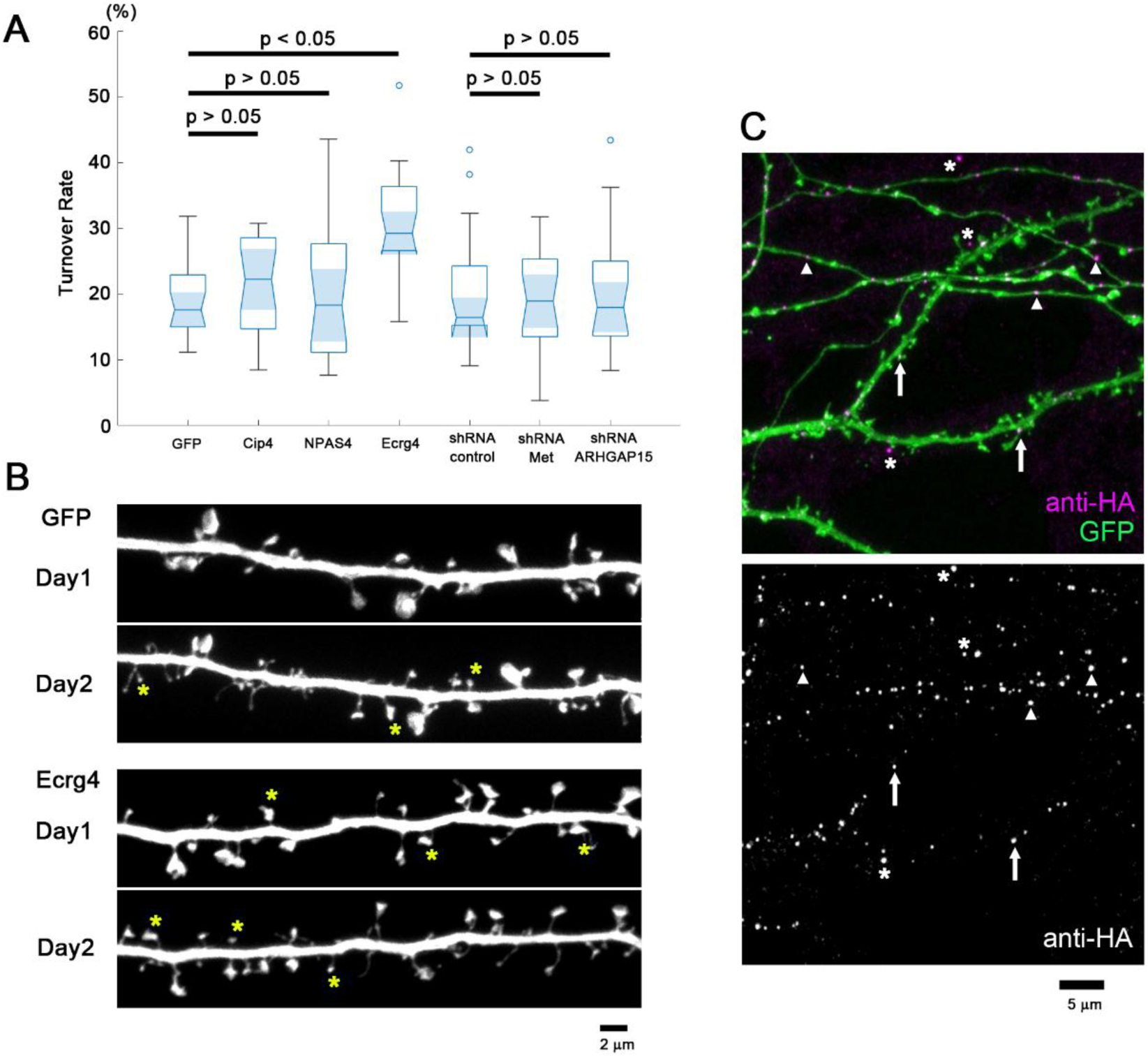
Manipulation of candidate gene expression in wild-type hippocampal neurons and its effects on spine turnover. (A) Turnover rates of dendritic spines in neurons transfected with an overexpression plasmid encoding Cip4, Npas4, or Ecrg4, or a plasmid encoding an shRNA targeting Met or Arhgap15, together with a GFP expression plasmid. Spine turnover rates were calculated from the images of GFP-expressing dendrites taken at an interval of 24 h. Among the five DEGs, only the upregulation of Ecrg4 selectively increased the spine turnover rate. Linear mixed-effects modeling with experiment as a random factor revealed a significant effect of overexpression treatment (F(3,8)=4.59, p=0.038), driven by Ecrg4 (p=0.013), whereas shRNA manipulations showed no significant effect (F(2,6)=0.29, p=0.76). (results from n = 22 dendrites/11 neurons/3 culture plates derived from independent primary culture for all conditions except the Met shRNA groups, where n = 21 dendrites from 11 neurons were included in the analysis). (B) Fluorescence images of dendritic segments expressing GFP or GFP plus Ecrg4 on days 1 and 2. Newly formed spines are marked by asterisks. Bar, 2 μm. (C) Images of dendrites and axons expressing HA-tagged Ecrg4 together with GFP. Anti-HA immunocytochemistry revealed the presence of immunopositive puncta both in the dendrites (arrows) and axons (arrowheads). Some clusters could be detected in the extracellular space (asterisks). The upper image is the overlay of the anti-HA signal (magenta) and GFP (green). The lower image shows the distribution of the anti-HA signals. Bar, 5 μm.

Ecrg4 encodes a precursor protein that yields several short hormone-like peptides via multiple cleavage sites^55^. Ecrg4 has been reported to function in cell proliferation, inflammation, and tumor suppression. However, there is limited information on the neuronal functions of Ecrg4. Therefore, we further analyzed the cellular distribution of Ecrg4 in wild-type hippocampal neurons to assess its potential synaptic functions. The Ecrg4 protein was detected in the membrane fraction of cultured hippocampal neurons (Supplementary Figure 10A). In addition, HA-tagged Ecrg4 showed punctate localization patterns within the neuronal cytoplasm (Figure 7C). A fraction of HA-tagged Ecrg4 puncta localized at axonal varicosities (arrowheads) or dendritic spines (arrows), together with some extracellular clusters (asterisks). HA-tagged Ecrg4 also colocalized with GFP-tagged neuropeptide Y (NPY) (Supplementary Figure 10B and C), suggesting transport in dense-core vesicles. Surface labeling of live cells also revealed clusters of Ecrg4 on axonal boutons and dendritic spines (Supplementary Figure 10D and E), supporting active exocytosis and tethering via its binding partners to the plasma membrane. Exogenous application of the synthetic Ecrg4 peptide enhanced spine turnover (Supplementary Figure 10F and G). Collectively, these observations suggest that the intracellular machinery for transporting and releasing Ecrg4 may be actively involved in regulating synapse formation and maturation.

### Rescue of the synaptic phenotype in schizophrenia-associated mouse models by Ecrg4 suppression

The spatial localization of Ecrg4 protein in wild-type neurons and upregulated gene expression in the hippocampus of schizophrenia model mice strongly suggest a contribution to the observed spine abnormalities. To examine this possibility, we tested whether Ecrg4 suppression by Ecrg4-shRNA rescues the synaptic phenotype of neurons derived from schizophrenia-associated mouse models (Supplementary Figure 9H and I). We introduced a GFP expression plasmid along with the Ecrg4-shRNA plasmid or the control shRNA plasmid into neurons derived from 22q11.2^del/+^ and Setd1a^+/-^ mice (Supplementary Table 1). The spine density plots generated from GFP-expressing neurons differed from those generated from DiI-labeled neurons. This difference may be due to the properties of SIM images obtained from either cell-surface DiI labeling or cytoplasmic GFP expression. Nevertheless, the subtracted (differential) distribution patterns between mutant and wild-type spines under the control shRNA condition indicate the enrichment of small spines (Figure 8A and J). From these plots, we identified the area showing enrichment of mutant spines (Figure 8B and K) and that of wild-type spines (Figure 8C and L). The pattern of small spine enrichment in mutant neurons shown in Figure 8A and J was no longer present in the plots generated by the subtraction of the wild-type spines under control shRNA from the mutant spines under Ecrg4 shRNA, indicating the possible rescue of the phenotype (Figure 8D and M). Consistent with this interpretation, enrichment in both the mutant- and wild-type–dominant regions was attenuated (Figure 8E, F, N, and O).

**Figure 8.**
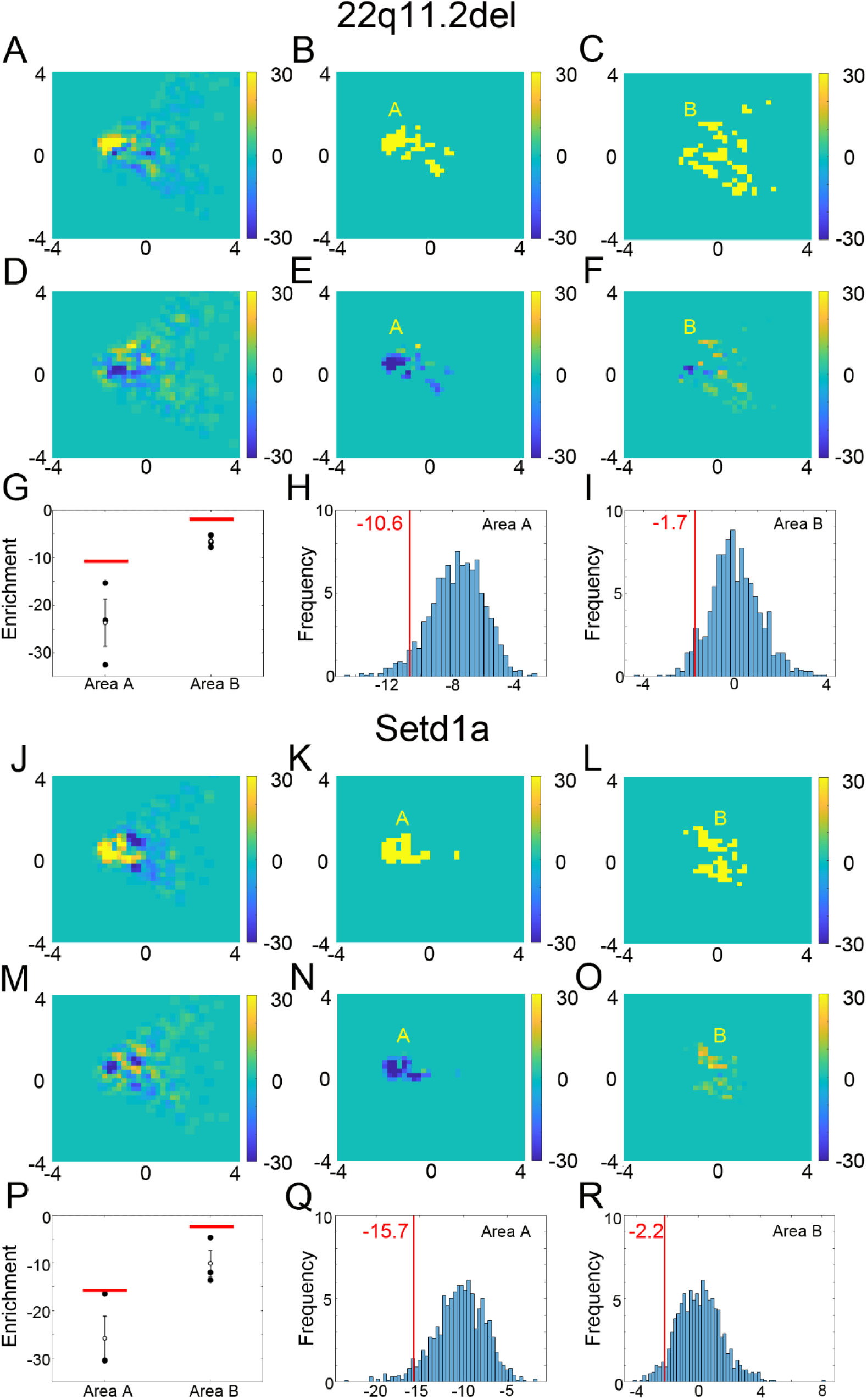
Altered spine population profiles after suppression of Ecrg4 expression in neurons derived from two schizophrenia-associated mouse models: the 22q11.2^del/+^ mouse model (A-I) and Setd1a^+/-^ mouse model (J-R). (A and J) Density plot obtained by subtracting wild-type spines from mutant spines. Both wild-type and mutant neurons expressed control shRNA. (B and K) Area enriched with the mutant spines under the control shRNA condition. (C and L) Area depleted of the mutant spines under the control shRNA condition. (D and M) Density plot obtained by subtraction of the wild-type spines under control shRNA from the mutant spines under Ecrg4 shRNA. This subtraction illustrates the normalizing effect of Ecrg4 shRNA. (E and N) Reduction of mutant spines in area A by Ecrg4 shRNA. The density plot, obtained by subtracting the mutant spines under control shRNA from those under Ecrg4 shRNA, shows the extent of normalization by Ecrg4 shRNA. (F and O) The increase of mutant spines in area B by Ecrg4 shRNA using the same subtraction method. (G and P) The extent of rescue by Ecrg4 shRNA in Area A and B for three independent culture experiments using either 22q11.2^del/+^ or Setd1a^+/-^ mouse models. The red lines show the values corresponding to statistical significance (p = 0.05), estimated from permutation analyses shown in (H, I, Q, and R). (H and Q) Permutation analysis for Area A to estimate the 95th percentile of the shuffled data. (I and R) Permutation analysis for Area B.

To determine whether the effect of Ecrg4 shRNA exceeded that expected from random variation in spine counts, we performed a permutation analysis. Spine identities were randomly reassigned between the two conditions while preserving the total number of spines in each dataset, and the differential density maps were recalculated for each permutation. This procedure generated a null distribution of enrichment values expected under random assignment (Figure 8H, I, Q, and R). The level of attenuation observed in both 22q11.2^del/+^ and Setd1a^+/-^ neurons exceeded the 95th percentile of the permuted distributions (p < 0.05), indicating that the obtained data were unlikely to be explained by stochastic variation alone (Figure 8G and P). This analysis supports the idea that Ecrg4 downregulation can rescue the spine-shape impairment observed in schizophrenia-related mouse models.

## Discussion

We developed an objective method for identifying population-level differences in spine nanostructures and applied it to cultured hippocampal neurons derived from multiple mouse models of psychiatric disorders. We identified two distinct groups of mouse models based on population-level spine properties. Notably, these two objectively identified groups correspond to mouse models with ASD-related gene mutations and schizophrenia-related gene mutations. A higher fraction of the small spine population was a common characteristic of schizophrenia-associated mouse models. Time-lapse imaging of spine dynamics revealed that this smaller spine phenotype is associated with a reduced initial size and growth of spines and their enhanced turnover. Screening for genes differentially expressed between mutant and control mice identified Ecrg4 overexpression as a potential molecular mechanism contributing to the schizophrenia spine phenotype. Indeed, suppression of Ecrg4 expression normalized the spine phenotype in neurons derived from schizophrenia-associated mouse models. These findings support the use of population-level analysis of spine nanostructures as a powerful approach to elucidating the heterogeneous synaptic impairments underlying psychiatric disorders.

Spine nanostructures provide information about developmental history, prior activity, and current functional properties, but these fine structural details were not accessible by conventional light microscopy. However, recent advances in super-resolution microscopy allow for the study of nanoscale spine morphology, including in living preparations, and have revealed new structural features related to important synaptic functions^11–13^. Another challenge to spine nanoscale imaging is the structural heterogeneity across large spine populations, which may obscure important population-level differences. Nonetheless, to understand the physiology and pathology of neural networks, it is essential to obtain comprehensive information about a large number of spines within a single neuron and across local neuronal networks. We previously reported a dimensional reduction technique coupled with machine learning for evaluating spine properties at the population level ^12^, and have further refined this method for the current investigation to extract population-level nanostructural features of spines in multiple mouse models of psychiatric disorders. This method enabled us to compare spine phenotypes across different mouse models objectively and group disease models based on similarities and differences in the spine population. Notably, the two identified groups corresponded to ASD- and schizophrenia-related gene mutations, supporting the idea that population-level analyses of spine nanostructures provide meaningful information on synaptic abnormalities and dysfunction.

Population-level spine properties were more homogeneous in schizophrenia models (those with gene mutations implicated in schizophrenia) than in the other 4 models studied, in part due to a shared tendency for smaller spines. This observation is consistent with previous studies on Setd1a mutant mice, which showed reduced spine width, decreased mushroom-type spines, and lower spine density in the prefrontal cortex^43,56,57^. In contrast to these findings, several previous studies reported reduced numbers of small spines in the postmortem cortical tissues of schizophrenia patients^22,58^. However, this discrepancy may be attributed to the developmental period, as we examined fetal neuronal spines after 3 weeks of in vitro differentiation, whereas the aforementioned studies included postmortem samples from patients over 40 years of age. One possible scenario that may help unify these observations is that spine density is reduced in the early postnatal period, and that this reduction triggers homeostatic plasticity, which, in turn, reduces the fraction of small spines in later life. Another potential explanation is that schizophrenia differentially affects regional spine properties (e.g., hippocampus versus cortex). Previous *in vivo* two-photon imaging studies reported higher spine turnover in the hippocampus than the cortex^59,60^. Therefore, the spine phenotype identified in this study may be specific to hippocampal pyramidal neurons. Finally, while the spine phenotype identified in the human postmortem brain undoubtedly resulted from complex interactions among genetic background, environmental influences, and regulation by non-neuronal cells, data from pure neuronal cultures are more likely to reflect the direct effects of schizophrenia-related gene mutations on synaptic functions. This property may be advantageous for identifying synaptic molecules that regulate synapse phenotypes in schizophrenia-related mouse models. However, the phenotype observed in the culture system requires confirmation using in vivo experiments of mouse models or human tissue samples. Efficient in vitro screening combined with reliable in vivo evaluation of synapses will facilitate translational research on mental disorders.

In addition to Ecrg4, these analyses identified several other DEGs common to mice harboring schizophrenia-related mutations. However, time-lapse imaging identified Ecrg4 as the most effective spine regulator, and we also found that Ecrg4 suppression reversed the abnormal spine phenotype in neurons derived from schizophrenia model mice. Ecrg4 encodes a precursor protein of hormone-like peptides ^61^. Exogenously expressed Ecrg4 exhibited a punctate pattern in both axonal and dendritic compartments, with a fraction of clusters present on the cell surface. This observation indicates that Ecrg4 may be secreted locally by axons or dendrites, thereby influencing synapses and adjacent neurons. Several signaling pathways act downstream of Ecrg4, including the stress-related NF-κB pathway and PI3K/Akt/mTOR pathway^55^. Ecrg4-derived peptides may also act through Toll-like receptor 4 (TLR4) or multiple scavenger receptors, such as LOX1^61^. While Ecrg4-related signaling pathways have been studied primarily in the context of inflammation and infection, there are few studies on Ecrg4-related signaling pathways in neurons. However, an Ecrg4-deficient mouse model demonstrated enhancements in neural stem cell proliferation and spatial learning, suggesting that overexpression may disrupt hippocampal function^54^. We suggest that Ecrg4 may be a critical regulator of neural development and synaptogenesis in the hippocampus. Further studies are warranted to identify Ecrg4 receptors and downstream signaling pathways and to investigate the potential dysfunction of these pathways in schizophrenia-associated mouse models.

The nanoscale features of dendritic spines in mouse models of Nlgn3^R451C/(y^ ^or^ ^R451C)^, Syngap1^+/−^, POGZ^Q1038R/+^, and 15q11-13^dup/+^, which we classified as being related to ASD, are highly heterogeneous. This heterogeneity may reflect the broad clinical spectrum of ASD, which ranges from mild impairments in social skills to severe intellectual disability. Accordingly, these four mouse models may represent distinct subgroups characterized by different degrees or forms of hippocampal dysfunction. Notably, among the ASD-related models, 15q11-13^dup/+^ showed population-level spine properties closer to those found in the 22q11.2^del/+^ and Setd1a^+/-^ mouse models. Although we classified 22q11.2^del/+^ and Setd1a^+/-^ as schizophrenia-related models, both 22q11.2 deletion syndrome and Setd1a haploinsufficiency in humans are also associated with ASD, suggesting substantial overlap in the genetic risk factors underlying ASD and schizophrenia. Further systematic analyses linking rare genetic variants to synaptic phenotypes in mouse models may provide important insights into the mechanisms underlying both shared and disorder-specific synaptic alterations in neurodevelopmental and psychiatric disorders.

The most important advantage of our analytical method is reproducibility, despite the substantial heterogeneity in genotype and phenotype among these mouse models. Thus, we are confident that the spine population analysis described here will help researchers in different laboratories obtain reliable and comprehensive information on spine properties. Further, this method could facilitate comparisons across a large number of mouse models harboring mutations associated with mental disorders. Spine morphology as measured in a reduced culture system can also demonstrate direct effects of gene mutations on neuronal phenotypes in the absence of indirect influences from nonneuronal cells or specific environments. Ultimately, these studies may identify the shared and specific pathogenic mechanisms of psychiatric disorders and lead to improved diagnosis and treatments.

## Materials and Methods

### Neuronal culture and genotyping of tissue samples from genetically modified mice

The following mouse models of psychiatric disorders were examined for spine nanostructural characteristics: male hemizygous or female homozygous neuroligin-3 R451C mutant (Nlgn3^R451C/(y^ ^or^ ^R451C)^), heterozygous Syngap1 mutant mice (Syngap1^+/−^), heterozygous POGZ mutant mice (POGZ^Q1038R/+^), mice with duplication of chromosome 7C (15q11-13^dup/+^; corresponding to human 15q11-13 duplication), mice heterozygous for deletion on chromosome 16B2,3 (3q29^del/+^; corresponding to human 3q29 deletion), mice heterozygous for deletion on chromosome 16qA13 (22q11.2^del/+^; corresponding to 1.5 Mb deletion on human 22q11.2), mice heterozygous for Setd1a mutantion (Setd1a^+/-^), and Ca^2+^/calmodulin-dependent protein kinase IIα kinase-dead mutant mice (CaMKIIα^K42R/K42R^). All animal experiments were reviewed and approved by the Institutional Animal Care and Use Committee of the University of Tokyo (permission number A2023M031-02).

For the production of embryos harboring heterozygous mutations, male heterozygous mutants were bred with wild-type female C57BL/6J or C57BL/6N mice (Japan SLC). Because neuronal cultures were prepared from wild-type and mutant littermates sharing the same genetic background, differences between the C57BL/6J and C57BL/6N sub-strains are unlikely to have affected the analysis. For the production of Nlgn3^R451C/(y^ ^or^ ^R451C)^ embryos, Nlgn3^R451C/y^ males and Nlgn3^R451C/+^ females were crossed. Embryos of CaMKIIα^K42R/K42R^ mice were obtained by crossing heterozygous CaMKIIα^K42R/+^ males and females. Both male and female embryos were used for the primary culture. Control cultures were prepared from littermate embryos without disease-related mutations on the background of either C57BL/6J or C57BL/6N. Genomic DNA was purified from E16 embryos prior to brain dissection using the QuickGene DNA tissue kit (WAKO), and genotypes were determined by PCR using either KapaTaq (Kapa Biosystems) or KOD FX Neo (TOYOBO) following the standard protocols provided by the manufacturers. The primer sequences used for genotyping were as follows:

Nlgn3^R451C/(y or R451C)^ genotyping:

Sense primer: 5′- GGTCAGAGCTGTCATTGTCAC-3′

Antisense primer: 5’- TGTACCAGGAATGGGAAGCAG-3’

Syngap1^+/-^ genotyping:

Sense primer for WT: 5’-GTCAGTGGGACATGGAAGTAG-3’

Sense primer for mutant: 5′-CTTCCTCGTGCTTTACGGTATC-3′

Antisense primer (common): 5’-CTGATCAGCCTGTCAGCAATG-3’

POGZ^Q1038R/+^genotyping:

Sense primer for WT: 5’- TCTGTGAAGAAGCTTCGGGTAGTAC-3’

Antisense primer for WT: 5′-GTCTCCTCATTTACAGGGAGCT-3′

Sense primer for mutant: 5’- GCAGCGGCTCCCCGTTAAC-3’

Antisense primer for mutant: 5′- AGCGCACAGCCCACTCATAG-3′

15q11-13^dup/+^ genotyping:

Sense primer: 5’- AGAGGAGGGCCTTACTAATTACTTA-3’

Antisense primer: 5′-ATATGTACTTTTGCATATAGTATAC -3′ 3q29^del/+^ genotyping:

Sense primer for WT: 5’- TTGGCACCACTCGCCCAAGTTATATCCACC-3’

Sense primer for mutant: 5’- CAAGGGGGAGGATTGGGAAGACAATAGCAG-3’

Antisense primer (common): 5’- GGTCATGCAAATTCTAGCAGTGAGTCATGAC-3’

22q11.2^del/+^ genotyping:

Sense primer for WT: 5’- GAGAAAGTGGGTGGGAAGGC -3’

Antisense primer for WT: 5’- GTCCCTCGCCACAGTCATAA -3’

Sense primer for mutant: 5’- CTAAGGAATGGTTCCGGCCA -3’

Antisense primer for mutant: 5′- TTTCACGGAGGCGGTATTCA-3′

Setd1a^+/-^ genotyping:

Sense primer: 5′-CTCGCCGCCATTTCTCTACATC-3′

Antisense primer: 5′-GTTCTGGAGGTTCTGGAGGTG-3′

CaMKIIα^K42R/K42R^ genotyping:

Sense primer: 5’-GGTCTTGAAGACTGTCTGGTGTGAGA-3’

Antisense primer: 5′-CACAGGCCAGTTTAGGTCTTGCTAGG-3′

After genotyping, primary hippocampal neuron cultures were prepared from mutant and corresponding control embryonic samples as described previously^12,62^. In brief, E16 hippocampi were dissected, minced, treated with trypsin (Gibco) and DNase (SIGMA), and dissociated mechanically into a cell suspension. Cells were resuspended in minimum essential medium supplemented with B18 supplement, L-glutamine (Gibco), and 5% fetal calf serum (Equitech-Bio), and plated onto glass-bottom dishes (MatTek, #1.5) precoated with poly-L-lysine (SIGMA). Glial proliferation was prevented by adding 5 μM Ara-C (SIGMA) 2 days after plating. For the comparison of spine nanostructure between wild-type and mutant neurons, culture samples were processed for imaging at 18-22 DIV. For comparison of 8 mutant mouse models using SIM imaging, all imaging data were obtained from the samples fixed at 19 DIV. For pharmacological treatment, 10 μM CNQX or 20 μM bicuculline was applied to the culture medium at 13 DIV, and samples were processed for imaging at 19 DIV.

### Transfection and fluorescence labeling of primary hippocampal neurons

Primary neurons were transfected with the indicated vectors using the calcium phosphate precipitation method at 8–9 DIV according to a previously described procedure^63^. All fluorescent proteins were expressed under the control of the β-actin promoter.

The expression of the shRNA constructs was induced using the pSilencer vector system. Briefly, the 19-nt target sequence and the reverse complement sequence of the target were ligated into the shRNA expression vector separated by a 9-nt spacer loop (TCTCTTGAA) (pSilencer 2.0-U6; Invitrogen). Stealth RNAi Negative Control Duplexes (Invitrogen) were used as control shRNAs. The targeting sequences for mouse Met, Arhgap15, and Ecrg4 are as follows:

Met RNAi sequence:

5’-GCAGTGAATTAGTTCGCTA-3

Arhgap15 RNAi sequence:

5’-AAGACAGATGTGAACATAC-3

Ecrg4 RNAi sequence:

5’- GAGGCTAAATTTGAAGAT-3’

The suppression of target protein expression by these transfected shRNAs was tested by immunoblotting lysates of COS-7 cells co-transfected with a target gene construct fused to a sequence encoding amino acids 98–106 of human influenza hemagglutinin (HA), all under the control of the β-actin promoter. Overexpression of Cip4, Ecrg4, and Npas4, genes upregulated in cultures derived from mouse models of schizophrenia, was induced by transfection of plasmids containing the corresponding HA-fused cDNA under the control of the β-actin promoter. Briefly, mouse Trip10 transcript variant 4 (Cip4; NM_001242391) and mouse Ecrg4 (NM_024283) were cloned, and an HA tag was inserted into the C-termini. Mouse Npas4 (NM_153553) was cloned according to a previous report^64^ and fused with a Flag-HA tag at its C-terminus. Human neuropeptide Y (NPY) tagged with green fluorescent protein (GFP, a kind gift from Dr. J. Takaska, NIH) was expressed under the control of the β-actin promoter.

For cell surface labeling of individual neurons in culture, cells were fixed and stained with 1,1’-dioctadecyl-3,3,3’,3’-tetramethylindocarbocyanine perchlorate (DiI; Molecular Probes) as described previously^12^.

### Immunocytochemistry

Cultured hippocampal neurons expressing HA-tagged constructs were fixed with 2% paraformaldehyde in PBS, permeabilized with 0.2% Triton X-100 in PBS, blocked with 5% normal goat serum, and reacted with rabbit anti-HA antibody (1:500; MBL), mouse anti-HA antibody (1:100; Biolegend), mouse anti-β-galactosidase antibody (1:500, SIGMA), rabbit anti-Ecrg4 (1:100, SIGMA), mouse anti-GM130 antibody (1:100, BD Biosciences), rabbit anti-Cip4/TRIP10 antibody (1:100, Proteintech), rabbit anti-NPAS4 antibody (1:100, Activity Signaling), and chicken anti-MAP2 antibody (1:500, PhosphoSolutions). After washing with PBS, samples were reacted with Alexa Fluor 546-labeled goat anti-rabbit IgG antibody (1:500; Invitrogen) and Alexa Fluor 647-labeled goat anti-mouse IgG antibody or anti-chicken IgY antibodies (1:500; Invitrogen). For cell surface labeling with GFP, neurons expressing Ecrg4-HA-SEP (superecliptic pHluorin) were reacted with anti-GFP VHH/nanobody conjugated to Alexa Fluor 647 (1:4,000; Chromotek) in culture medium at 37°C.

### Structured illumination microscopy (SIM) imaging

For 3D-SIM imaging, we used an inverted microscope (ECLIPSE Ti-E, NIKON) equipped with an N-SIM-E system, 405, 488, 561, and 640 nm diode lasers (LU-NV, NIKON), and an oil-immersion TIRF objective lens (SR Apo TIRF 100 ×, N.A. 1.49, NIKON). All SIM images were acquired under strictly controlled conditions, including a stage temperature of 28–29°C, to minimize the position and aberration fluctuations. Spherical aberrations induced by refractive index mismatch were reduced by manually adjusting the objective lens correction collar. Images were acquired using an EMCCD camera (iXon3 DU-897E, Andor Technology) with 512 × 512 pixels (each pixel, 16 μm square) operated in the read-out mode at 1 MHz with 16-bit analog-to-digital conversion of the EM gain. All image-processing steps were performed in three dimensions. The acquired image datasets were computationally reconstructed using the reconstruction stack algorithm V2.10 of NIS-Elements AR (NIKON). An image stack consisted of 63 axial (z) planes of 512 × 512 pixels in the x- and y-dimensions, with a pixel size of 32 × 32 nm. All 3D images were reconstructed in 120-nm z-steps spanning 7.56 μm in the z-axis. The voxel size met the Nyquist criterion. Only dendritic segments spatially isolated from other fluorescent objects were selected for analysis because objects brighter than the targeted dendritic segments made image thresholding unreliable. Quality checks of the acquired SIM images were performed as described previously using the SIMcheck plugin for ImageJ^65^.

### Spine isolation and shape measurement

Spines were isolated by automated image thresholding using a previously described method^12^. Briefly, the reconstructed SIM image stacks were first processed by multilevel thresholding using the MATLAB built-in function [multithresh()] to produce binary images, which were further processed by geodesic active contours using the MATLAB built-in function [activecontour()]. The binary image stacks generated by thresholding the SIM images were then processed for the automated detection of spines. First, the dendritic shafts were fitted with elliptic cylinders and voxel clusters outside the best-fit elliptic cylinders were identified as spine candidates. Next, the spine candidates were classified by volume and shape. After selection, the junction between the dendritic shaft and spine was determined. Finally, the nanostructural parameters of isolated spines were measured. These steps are basically the same as those in our previous algorithm, but several points were improved, as described in the following sections (The original code for SIM image processing is available at https://shigeookabe.github.io/download-page-SIM/ with a password of “simspineimage”).

(1) Dendritic shaft fitting

We first estimated the shaft volume using the custom-made function “SIM_Fitdendrite” as previously described. In the new algorithm, the candidate spines were subsequently evaluated to determine whether their bases were large and spread on the shaft surface. If the spine base was large, the basal voxels were removed using the function “SIM_Removevoxel.”

(2) Selection of the spines

Using the custom-made function “SIM_Position,” spines with volumes too small, too elongated, or located at the image edge were removed as previously described. Small spine candidates with volumes less than twice the volume of a rectangular cuboid with edge lengths equal to the theoretical resolutions of SIM images (<0.01 μm^3^) were rejected.

(3) Adjustment of the spine–shaft junction

We determined the location of spine–shaft junction using the custom-made function “SIM_Junction.” This step was identical to our previous algorithm. Next, spines with more than one connection site to the shaft were detected using the function “SIM_Neck” and eliminated from further analysis.

(4) Measurement of nanostructural parameters

First, the angle of the spines was adjusted to be perpendicular to the long axis of the dendritic shaft and within the xy-plane using the function “SIM_Angle.” Next, nanostructural parameters were measured using the function “SIM_Shape.” In our previous spine shape measurement study, a polygon mesh-based calculation was performed. In the current study, we took a different approach using voxel-based measurement. The calculation of structural parameters using voxel data was more straightforward and required fewer calculation steps than the method based on polygon mesh data. Spines were divided into 160-nm segments along the long axis using the function “SIM_Shape.” These spine segments were used to calculate the following 64 nanostructural parameters: spine length, surface area, base surface area, total volume, volume of each spine segment (20 segments), convex hull volume of each spine segment (20 segments), and convex hull ratio of each spine segment (20 segments).

### Analysis of the spine population data

Differences in spine phenotype among model mice were analyzed at the population level by PCA. We selected five parameters (spine length, base surface, total volume, and volumes of the fifth and fifteenth spine segments) for the PCA-based dimensional reduction. Two parameters that reflect the principal structural features (length and volume) were first selected. Second, three other parameters (base surface, volumes of the fifth and fifteenth spine segments) that were mutually independent and also independent from the first two parameters were chosen (pairwise correlation coefficients < 0.3). The initial 64 parameters included the volume of each spine segment (20 segments), the convex hull volume of each spine segment (20 segments), and the convex hull ratio of each spine segment (20 segments). These parameters showed higher correlation coefficients but did not improve PCA performance. We also confirmed that PCA using all 64 parameters yields a cross-correlation map similar to that shown in Fig. 2B. The first three principal components (PC1–PC3) covered approximately 90% of the variance in the data.

The 2D presentation of spine population data using PC1 and PC2 reflected the overall pattern of the spine shape distribution, with thin, mushroom, and stubby spines separated in the feature space. We found that the overall pattern of spine population data projected to the PC1-PC2 plane differs between DiI-labeled neurons and GFP-expressing neurons. This difference may be due to the properties of SIM images obtained from either cell-surface labeling or cytoplasmic GFP expression. To facilitate the direct comparison of the spine population data in Figure 3 (DiI-labeled neurons) and Figure 8 (GFP-expressing neurons), we transformed the raw data of GFP- expressing neurons using the weight matrix calculated from the data for DiI-labeled neurons.

After PCA, relative spine densities (spine number per area in the feature space) were mapped onto an 80 × 80 grid spanning ±4 standard deviations along PC1 and PC2. Using this procedure, two density plots were generated: one from culture samples of disease model mice and the other from corresponding control cultures. The two density plots were then subtracted to extract the local differences in spine numbers. Because each grid contains spines with similar structural parameters, the subtracted plot reveals regions of mutant-dominant and wild-type-dominant spine distributions in the PC1–PC2 plane.

The similarity between the two subtracted density plots was evaluated by calculating the 2D cross-correlation C using the following equation

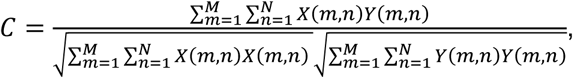

where X and Y correspond to the two subtracted density matrices. The 11 different subtracted density plots shown in Figure 2 were clustered using the MATLAB function “linkage,” and the average distance between all pairs of elements in the two samples was calculated using the cluster algorithm.

For Figure 3, the PC1–PC2 feature space was divided into four subareas with distinct shape properties: (1) small and short, (2) small and long, (3) large and short, and (4) large and long. The median spine volume and length were first calculated from the wild-type population, and spine samples were classified into four subgroups based on these median values. Each position of the 80 × 80 grid was then labeled according to the dominant subgroup of spines it contained, yielding a map of four subareas corresponding to the four spine shape categories.

In Figure 3, 8, and Supplementary Figure 4, areas in the feature space enriched with mutant or control spines were extracted by searching the 80 × 80 grid and identifying those where the difference in relative density was larger than the mean plus 3 × SD. Spine axial profiles in Supplementary Figure 6 were then analyzed within areas enriched with either mutant or control spines.

### Time-lapse imaging of dendritic spines and image analysis

Live cell 3D imaging was performed using an A1 confocal laser scanning microscope (NIKON) equipped with 405, 488, 561, and 640 nm diode lasers (LU-NV, NIKON). Cells at 15–18 DIV were maintained at 37°C under a continuous flow of 5% CO_2_ to maintain medium pH using a heater stage system (INUG2H-TIZSH, Tokai Hit) with a lid designed for a glass-bottom dish to minimize evaporation. Horizontal and vertical drifts were controlled by a motorized stage and a perfect focus system, respectively. Images were acquired with an oil immersion objective lens (60×, NA 1.4, NIKON) at a confocal aperture of 1 AU. The imaged volume was 30.7 × 30.7 × 7.56 μm in x, y, and z directions, respectively, and image stacks consisted of 17 planes separated by 500 nm. Time-lapse images were recorded either at short intervals (15 min intervals for 24 h) or long intervals (two time points separated by 24 h). Cultured neurons imaged at a 24-h interval were then fixed and immunostained with anti-HA to confirm the expression levels of the transfected genes. The effect of exogenously applied Ecrg4 peptide (CΔ16; the C-terminal ECRG4 fragment of 16 aa, 5.4 μM, BEX CO., LTD.) was evaluated by taking time-lapse images at 24 h intervals, with the first image taken just before the peptide application.

The image stacks obtained by time-lapse imaging were analyzed using custom-made MATLAB scripts. First, 2D images were generated by maximal intensity projection of z-stack images. These 2D images were processed to transform the outlines of dendritic shafts to be aligned with straight lines. Spines were subsequently identified as objects extending beyond the straight edges of the dendritic shafts. Identified spines were segmented, and total fluorescence intensities were measured as a proxy for spine volume. We classified the spines as stable (present throughout the imaging period), newly formed (appearing within 24 h imaging period), and eliminated (lost within 24 h imaging period). Spine turnover rate was calculated as the sum of newly formed and eliminated spines during a 24-h interval, normalized to the total number of spines present at the beginning and end of the interval. The resulting value was expressed as a percentage. For newly formed and eliminated spines, we generated temporal profiles of growth and shrinkage using the total fluorescence intensity values.

### RNA sequencing and gene expression analysis

Total RNA was isolated from cultured hippocampal neurons at 19 DIV using the RNeasy Mini Kit (QIAGEN) with the RNase-Free DNase Set (QIAGEN) according to the manufacturer’s protocols. The quality of the extracted RNA was initially assessed using a NanoDrop spectrophotometer, and the RNA integrity number (RIN) score was calculated for each sample using a Bioanalyzer (Agilent Technologies). The yields were quantified using a Qubit Fluorometer (Thermo Fisher Scientific). Sequencing libraries were prepared using the NEBNext Ultra Directional RNA Library Prep Kit for directional libraries (New England BioLabs) and the KAPA HTP Library Preparation Kit (KAPA Biosystems) according to the manufacturers’ protocols. Paired-end RNA-seq libraries were sequenced using the Illumina HiSeq and MGI DNBSEQ-G400 platforms. Three independent culture preparations were used for RNA sequencing of each mouse model and control library. Following quality control, the reads were aligned with the mouse reference genome GRCm38/mm10 (GenBank assembly ID: GCA_000001635.2). Expression levels of genes were analyzed based on the transcripts per million and differential expression analysis was conducted using the DESeq2 package in R^66^. Differentially expressed genes (DEGs) were selected according to an adjusted P < 0.05 and a (log2)-fold change greater than 0.5.

### Simulation of spine development

Spine growth was simulated using a dendrite model consisting of 1,000 segments, each 0.25 μm in length (250 μm in total). The spine distribution was updated every 20 min over 10 days, covering the initial phase of spine development from 8 DIV to 17 DIV. The initial probability of new spine generation was set at 7.5 × 10^−3^ per segment per hour and decreased thereafter in proportion to the accrued spine density increase (by the final 24 h, the probability of spine generation [mean ± SD] was 3.2 × 10^−3^ ± 0.23 × 10^−3^/segment/h). This probability function yielded an average of 1.2 ± 0.22 new spines per 10 μm of dendrite every 24 h, in good agreement with the spine dynamics measured by time-lapse imaging from 15–18 DIV (1.3 ± 0.63 new spines per 10 μm of dendrite every 24 h). The model also set the increase in spine volume (V1) from 6.7 × 10^−3^ to 3.3 × 10^−2^ in 20 min. Spines in the early growth phase were treated as unstable, with loss probability P1 ranging from 9.0 × 10^−3^ to 0.1 every 20 min. Spines were modeled as stabilized when the individual volume reached a relative value of 1.0. These stabilized spines then entered a shrinking phase with a probability P2 ranging from 2.9 × 10^−3^ to 4.0 × 10^−3^ in 20 min. The shrinking rate V2 was set from 6.7 × 10^−3^ to 3.3 × 10^−2^ per 20 min. In simulations, the four parameters V1, V2, P1, and P2 were varied within these indicated ranges to find the combination producing the best fit with experimental data on the difference in spine lifetime distribution profile and turnover rate at 24 h (the actual measurement interval). The simulation with the best-fit parameters yielded a spine density comparable to measured data (0.57 ± 0.029 vs. 0.70 ± 0.17 spines/μm of dendrite).

### Western blotting

Total protein extracts from cultured hippocampal neurons were separated into cytosolic and membrane fractions using Trident Membrane Protein Extraction Kit (GeneTex) according to the manufacturer’s protocol and separated on 10% sodium dodecyl sulfate (SDS)-polyacrylamide gels or 15%–20% Tricine-SDS-polyacrylamide gels. The separated proteins were then transferred onto nitrocellulose membranes (Millipore) or PVDF membranes (Bio-Rad) using a wet transfer system. Membranes were blocked with 5% bovine serum albumin in Tris-buffered saline plus Tween-20 (TBS-T, 20 mM Tris-HCl, 137 mM NaCl, 0.2% Tween-20) for 1 h at room temperature, then incubated overnight at 4°C with rabbit anti-Ecrg4 (1:1,000, SIGMA), rabbit anti-GluA1 (1:1,000, Nittobo Medical), rabbit anti-GFP (1:3,000, MBL), rabbit anti-dsRed2 (1:4,000, Nittobo Medical), mouse anti-α-tubulin (1:3,000, SIGMA) as indicated. The membranes were washed and incubated with HRP-conjugated secondary antibody against mouse or rabbit IgG (1:10,000, Amersham) for 2 h. Signals from the target protein bands were detected using a SuperSignal™ West Femto or Atto detection kit (Pierce) and imaged for quantitation (densitometry) using a ChemiDoc imaging system (Bio-Rad). We isolated the fraction of synaptosomes using Syn-PER™ Synaptic Protein Extraction Reagent (Thermo Scientific) in Supplementary Figure 10A, according to the manufacture’s protocols.

### Statistics

The statistical tests used for each experiment and the sample sizes (spines, dendrites, cells, and plates) are indicated in the corresponding text or figure legends. Statistical significance was determined as follows:

Figure 3G-J: One-way ANOVA followed by Tukey’s post hoc test.

Figure 5A-C: A linear mixed-effects model with genotype as a fixed effect and plate, cell, and dendrite as nested random effects.

Figure 5D-F: A linear mixed-effects model with genotype as a fixed effect and plate, cell, dendrite, and spine as nested random effects.

Figure 5G-L: A linear mixed-effects model with genotype as a fixed effect and plate, cell, dendrite, spine, and time as nested random effects.

Figure 7A: A linear mixed-effects model with treatment as a fixed effect and plate, cell, and dendrite as nested random effects.

Figure 8: Three independent imaging datasets evaluated by permutation tests.

Statistical significance was set at P < 0.05.

## Acknowledgments

We thank Toru Takumi for the 15q11-13^dup/+^ mice, Katsuhiko Tabuchi for Nlgn3^R451C/(y^ ^or^ ^R451C)^ mice, Yoko Yamagata for CaMKIIα^K42R/K42R^ mice, and Justin Taraska for the NPY tagged with GFP. This work was supported by Grants-in-Aid for Scientific Research (25H01038, 20H00481, 20H05894, 20H05895 to S.O.), the Japan Agency for Medical Research and Development (JP19gm1310003 to S.O., T. N., and Y. G., JP22jm0210097, JP23wm062500 to S.O., JP19dm0207071 to A. A.), the Naito Foundation, and the Uehara Memorial Foundation.

## Author contributions

Y.K. and S.O. conceived of the project and designed the methodology. Y.K. and Q.L. performed the experiments. R. S., A.A. and T.N. provided the animal resources and provided advice on analysis. Y.G. conducted the gene expression analysis. S.O. and Y.K. wrote the manuscript, with contributions from A.A., T.N., and Y.G.

## Declarations of generative AI and AI-assisted technologies in the writing process

The authors utilized Grammarly (https://app.grammarly.com/) for editing the text. The authors take full responsibility for the contents of the manuscript.

## Resource availability

The original code for SIM image processing is available at https://shigeookabe.github.io/download-page-SIM/ with a password of “simspineimage”.

## Supplementary Figures and Legends

**Supplementary Figure 1.**
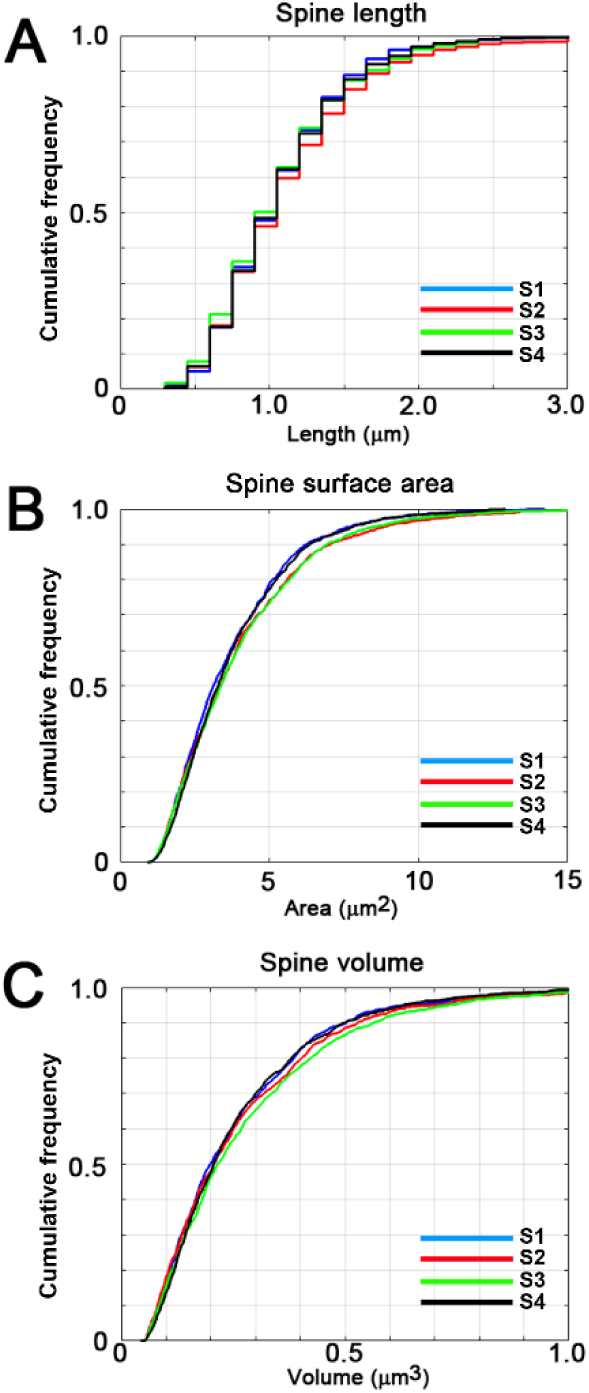
Cumulative frequency plots of spine length, surface area, and volume measured in four independent experiments performed > 2 months apart. The cumulative frequency distributions of spine length (A), spine surface area (B), and spine volume (C) were obtained from wild-type data acquired by SIM imaging across four mouse models: Nlgn3^R451C/(y^ ^or^ ^R451C)^ (S1), Syngap1^+/−^ (S2), POGZ^Q1038R/+^ (S3), and 15q11-13^dup/+^ (S4). The Kolmogorov–Smirnov test detected significant differences in only three of 18 possible pairwise comparisons: surface area of (S1) vs. (S3) (p = 0.017), volume of (S1) vs. (S3) (p = 0.032), and volume of (S3) vs. (S4) (p = 0.038).

**Supplementary Figure 2.**
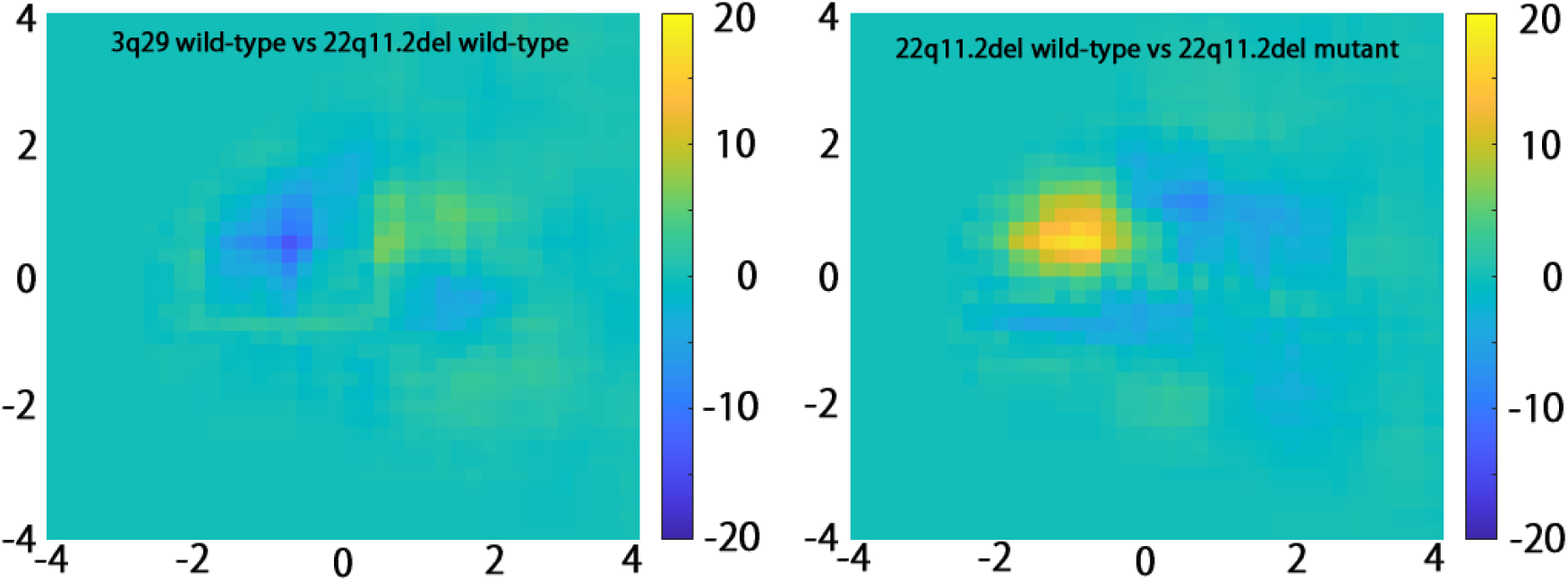
Subtracted density plots comparing wild-type cultures from 3q29^del/+^ and 22q11.2^del/+^ mice (left) and wild-type versus mutant cultures from the 22q11.2^del/+^ line (right). The differences observed between independent wild-type culture preparations were substantially smaller than those observed between wild-type and mutant cultures, indicating that genotype-dependent effects exceeded inter-preparation variability.

**Supplementary Figure 3.**
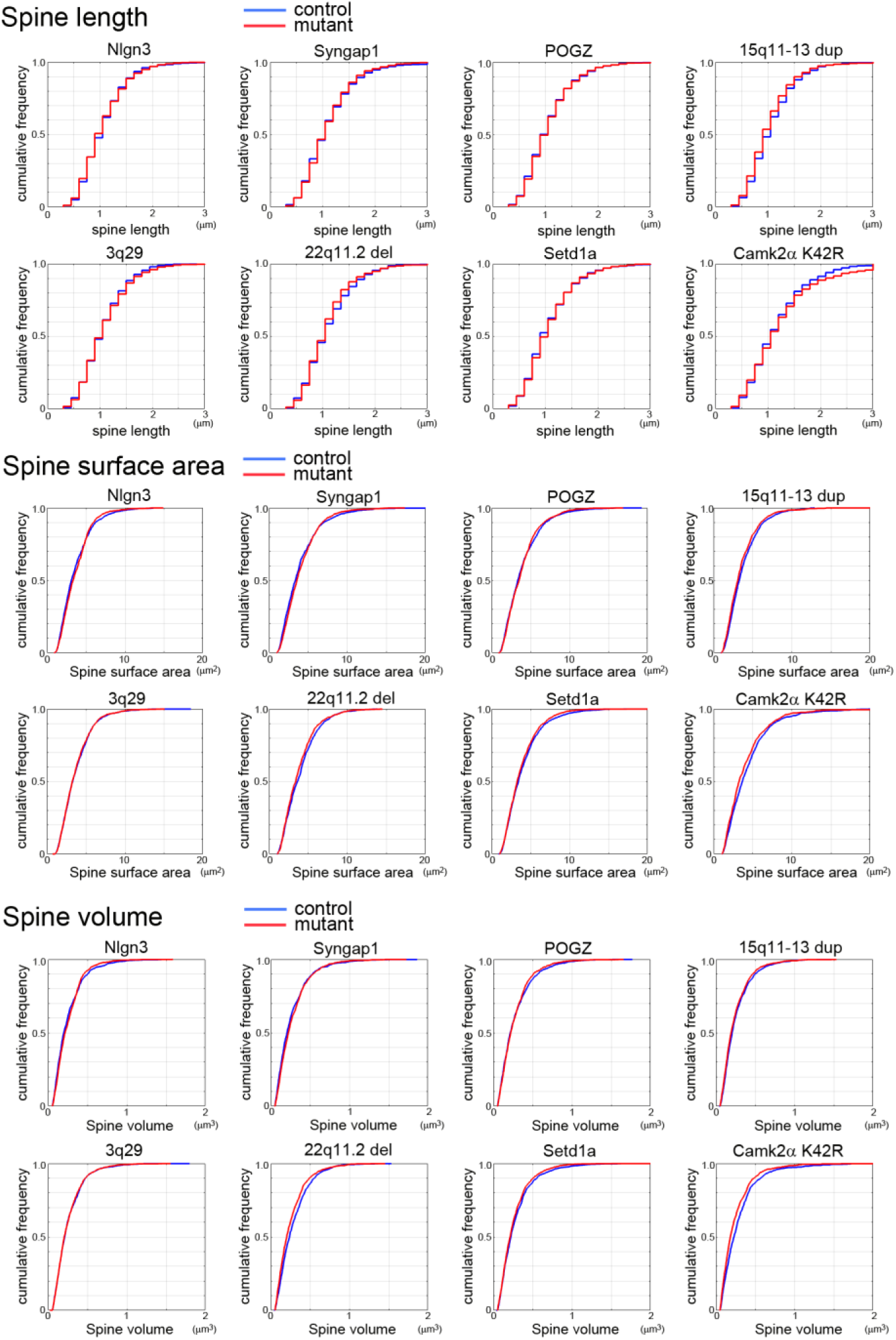
Cumulative frequency plots of spine length, surface area, and volume for the eight mouse mutants: Nlgn3^R451C/(y^ ^or^ ^R451C)^, Syngap1^+/−^, POGZ^Q1038R/+^, 15q11-13^dup/+^, 3q29^del/+^, 22q11.2^del/+^, Setd1a^+/-^, and CaMKIIα^K42R/K42R^.

**Supplementary Figure 4.**
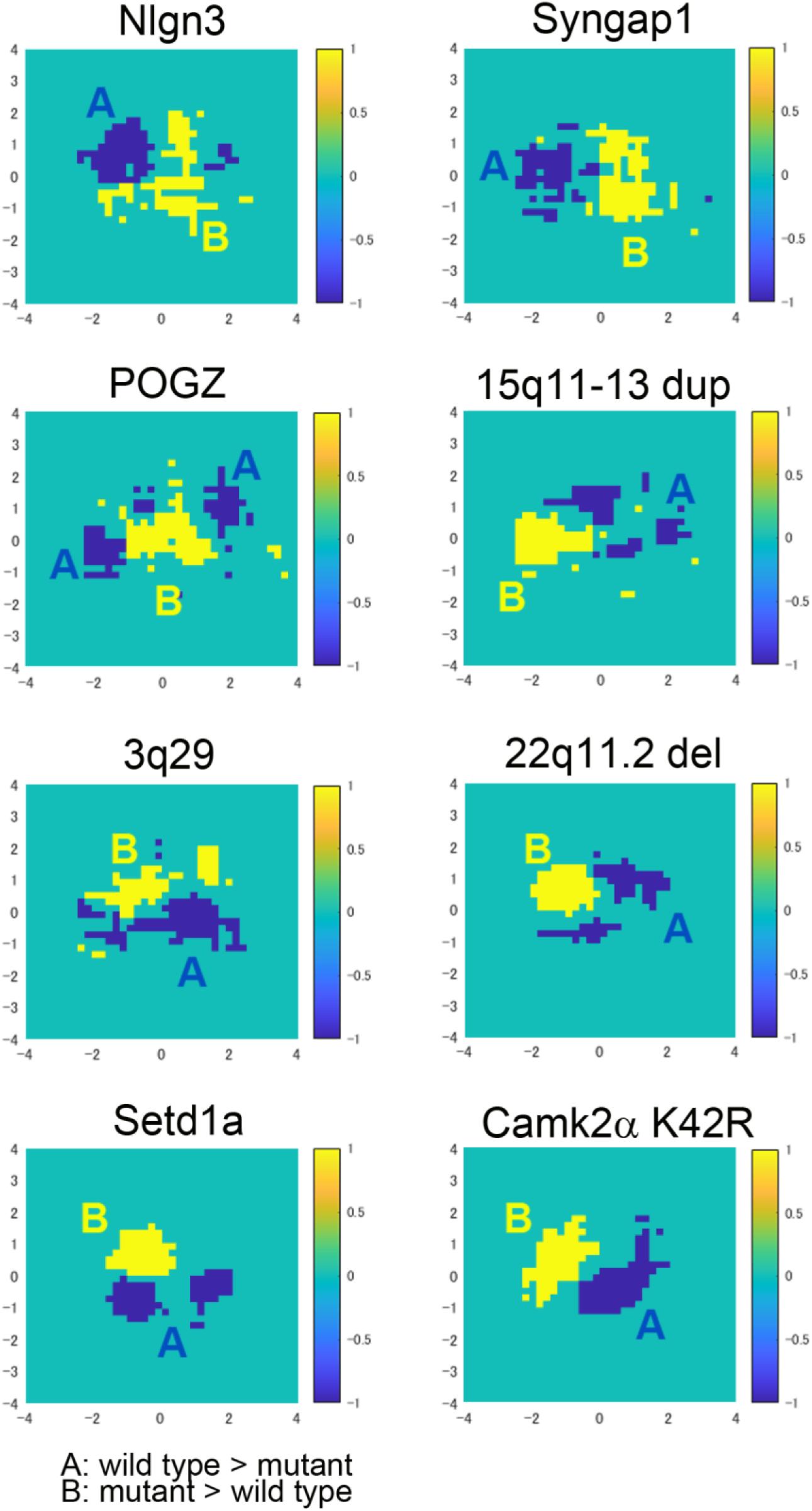
Areas where control (A: blue) or mutant (B: yellow) spines show a higher density within the feature space of the PC1-PC2 plane. The plots for all 8 mouse models are presented. The differences in population-level spine properties between ASD- and schizophrenia-associated mouse models were preserved within two groups. The areas enriched with mutant spines in Nlgn3^R451C/(y^ ^or^ ^R451C)^, Syngap1^+/−,^ and POGZ^Q1038R/+^ models were on the right side of the plot, suggesting the abundance of large spines in this group. In contrast, the areas enriched with mutant spines in 3q29^del/+^, 22q11.2^del/+^, Setd1a^+/-^, and CaMKIIα^K42R/K42R^ models were positioned on the left side of the plot, indicating the abundance of small spines.

**Supplementary Figure 5.**
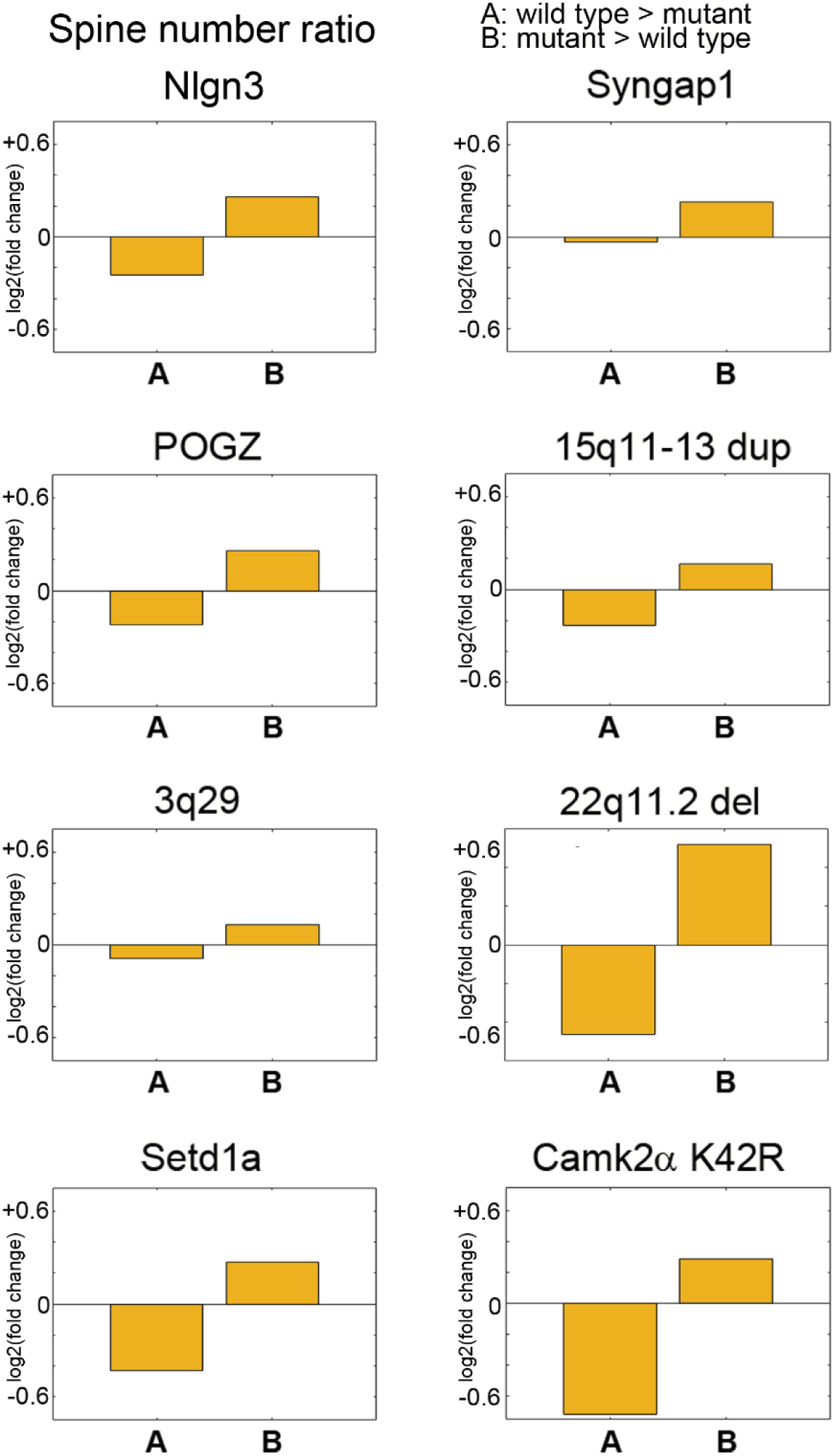
The relative numbers of spines within areas A and B in the feature space from Supplementary Figure 4. The densities of mutant spines were higher in area B than in area A.

**Supplementary Figure 6.**
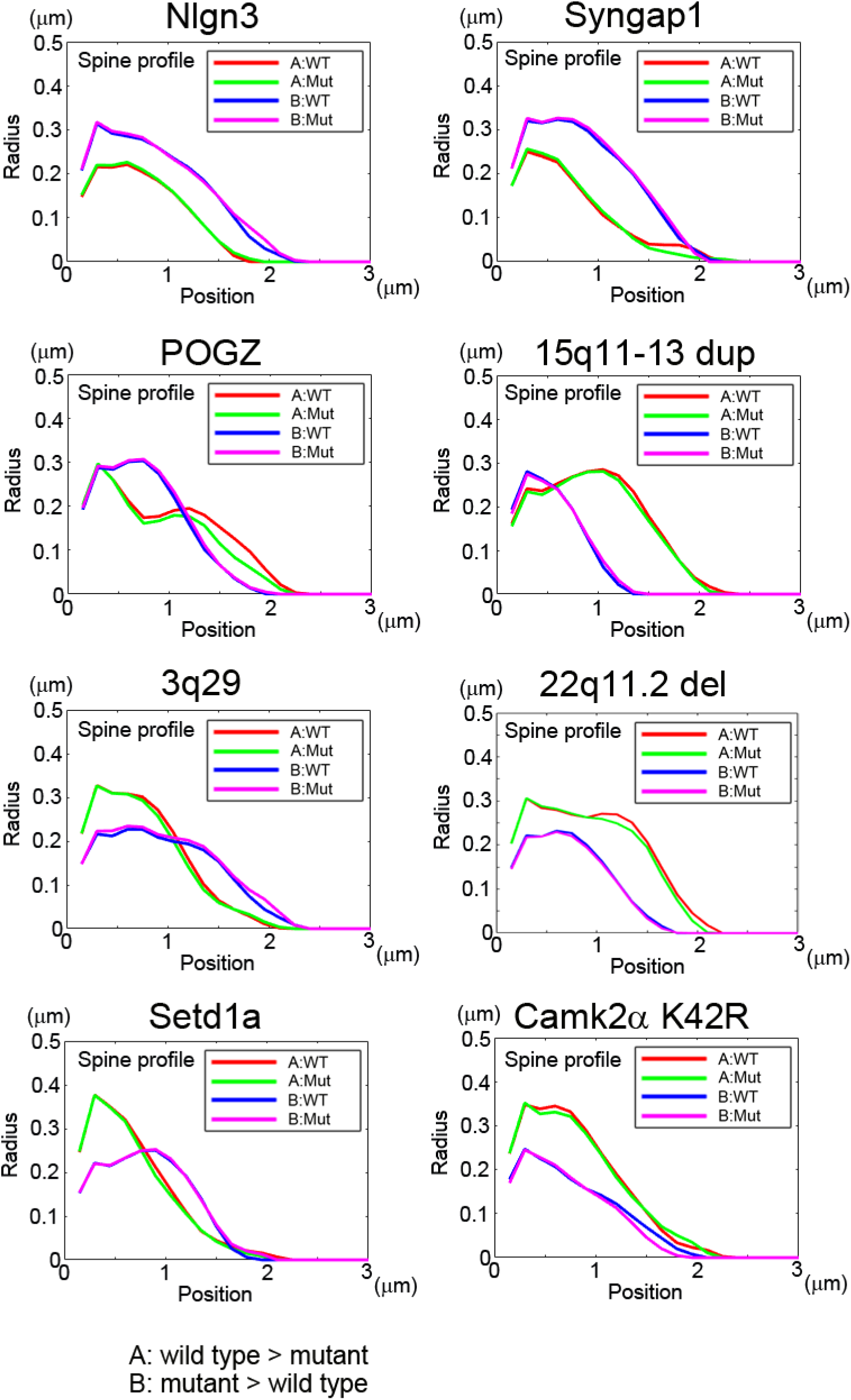
Profiles of different spine populations (spines in the control-dominant area A and the mutant-dominant area B for both wild-type and mutant neurons) were visualized by plotting the radius along the long axis. The numbers of control and mutant spines included in the analysis are as follows: Nlgn3^R451C/(y^ ^or^ ^R451C)^, n = 775 in area A and n = 513 in area B; Syngap1^+/−^, n = 668 in area A and n = 500 in area B; POGZ^Q1038R/+^, n = 580 in area A and n = 815 in area B; 15q11-13^dup/+^, n = 378 in area A and n = 797 in area B; 3q29^del/+^, n = 627 in area A and n = 824 in area B; 22q11.2^del/+^, n = 308 in area A and n = 575 in area B; Setd1a^+/-^, n = 472 in area A and n = 808 in area B; CaMKIIα^K42R/K42R^, n = 237 in area A and n = 634 in area B.

**Supplementary Figure 7.**
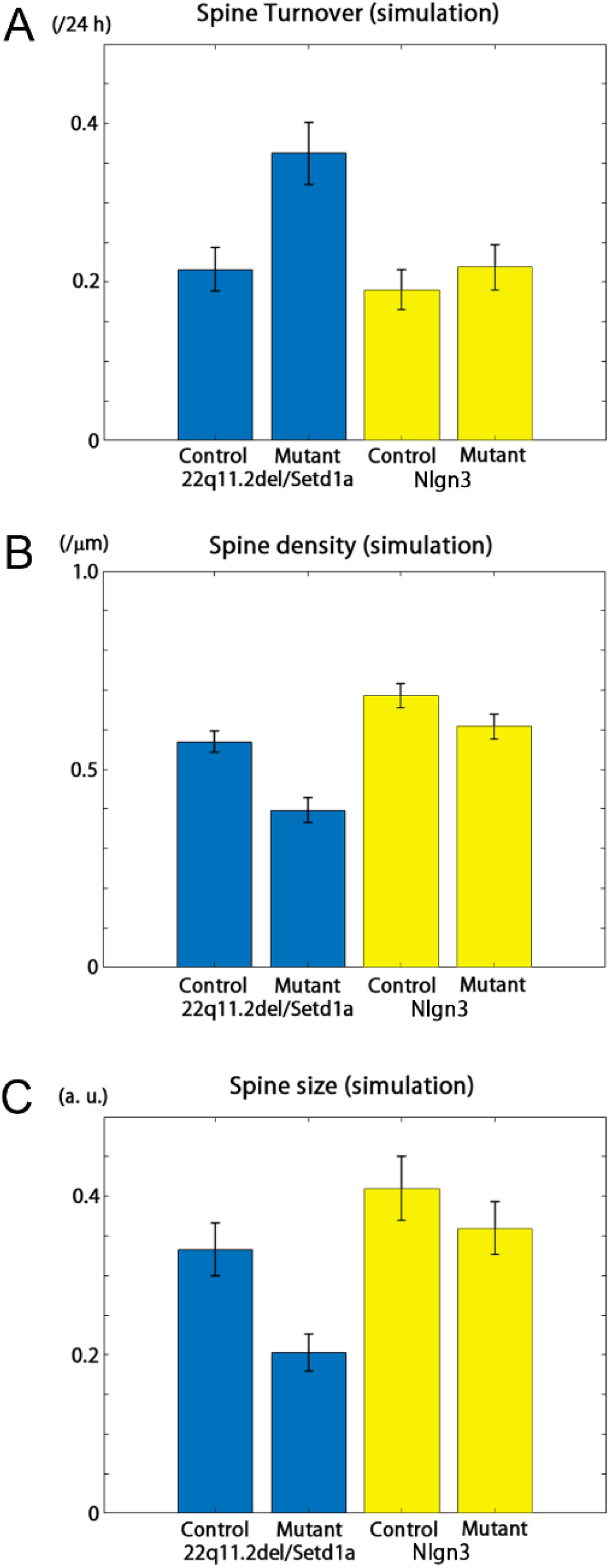
Spine turnover (A), density (B), and size (C) from simulation data. Simulation parameters were tuned to fit experimental results from control and mutant models. Means and standard deviations are shown.

**Supplementary Figure 8.**
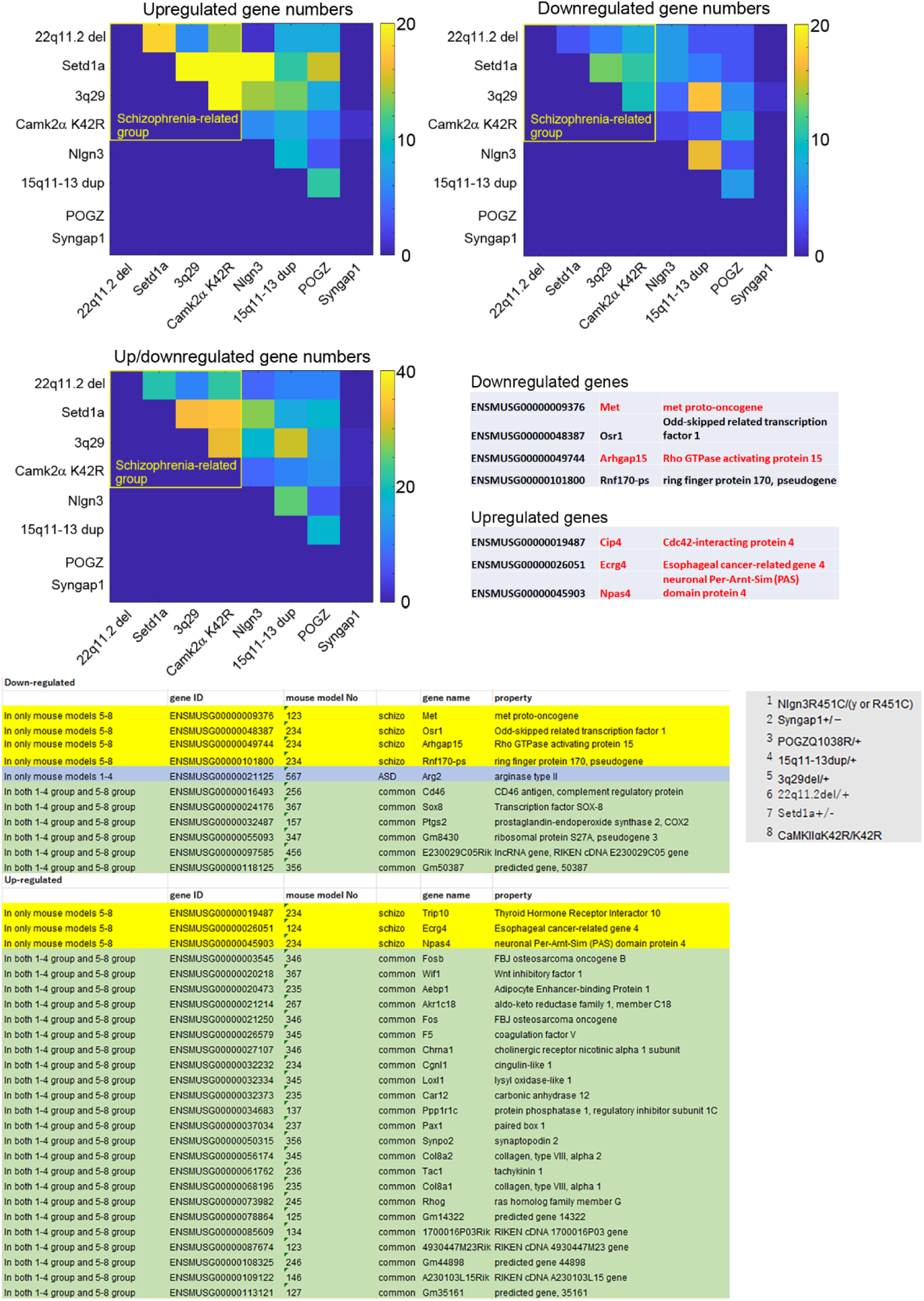
Pseudocolor presentation of differentially expressed genes (DEGs) between mutant and corresponding control mice. The number of shared DEGs was higher in the schizophrenia-related mouse models than in the ASD-related mouse models. Mouse gene identifiers (ENSMUSG) and gene names for DEGs shared by three or four schizophrenia-associated mouse models and not differentially expressed in ASD-associated mouse models are presented. DEGs analyzed for their effects on spines are shown in red. Color indicates the number of DEGs. The lower table shows all DEGs shared by three or four mouse models, irrespective of their grouping as ASD-related or schizophrenia-related.

**Supplementary Figure 9.**
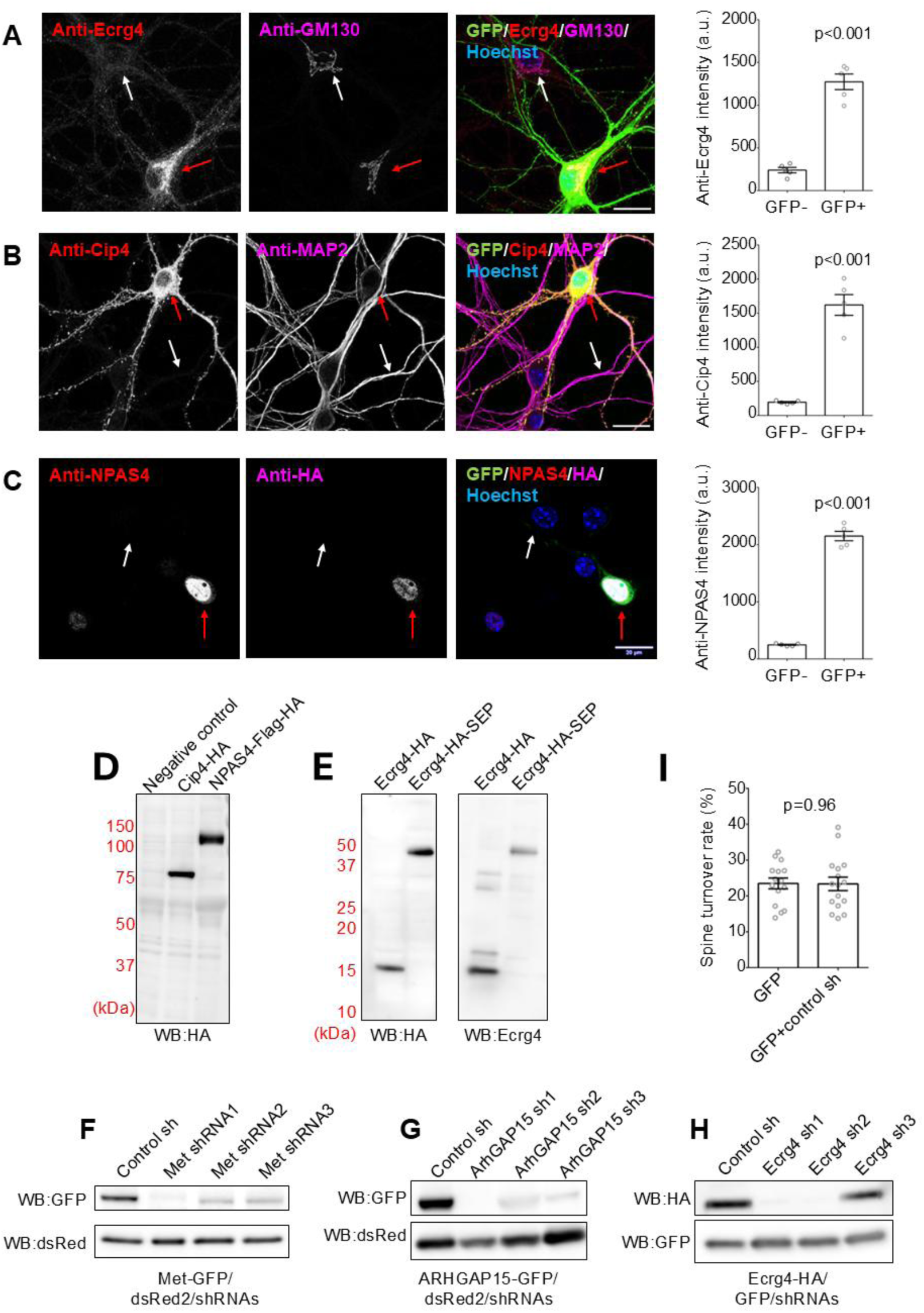
(A-C) Validation of overexpressed tagged constructs by immunocytochemistry in hippocampal neurons. Bar = 20 μm. The data are presented as mean ± SEM (N = 5 neurons each). Red allows; transfected neurons, white arrows; non-transfected neurons. (A) Immunocytochemistry of hippocampal neurons expressing Ecrg4-HA with GFP using anti-Ecrg4 antibody. Anti-GM130 antibody staining detected the Golgi apparatus. (B) Immunocytochemistry of hippocampal neurons expressing Cip4-HA with GFP using anti-Cip4 antibody. Anti-MAP2 antibody detected neuronal dendrites. (C) Anti-NPAS4 and anti-HA immunocytochemistry of hippocampal neurons expressing NPAS4-Flag-HA with GFP. (D-E) Immunoblot verification of tagged construct expression in COS-7 cells. (D) Immunoblotting in total cell lysate from COS-7 cells expressing Cip4-HA or NPAS4-Flag-HA. The estimated molecular weights of Cip4-HA and NPAS4-Flag-HA are 64 kDa and 89 kDa, respectively. (E) Immunoblotting using both anti-HA and anti-Ecrg4 in total cell lysate from COS-7 cells expressing Ecrg4-HA or Ecrg4-HA-SEP. The estimated molecular weights of Ecrg4, Ecrg4-HA, and Ecrg4-HA-SEP are 17 kDa, 18 kDa, and 45 kDa, respectively. (F-H) Verification of knock-down effects of shRNAs by immunoblotting of COS-7 cells. Immunoblotting in total cell lysates from COS-7 cells transfected with the expression vectors of either Met-GFP (F), ArhGAP15-GFP (G), or Ecrg4-HA (H), together with dsRed2 and shRNAs targeting the corresponding transcripts. (I) Measurement of spine turnover rate in neurons transfected with a control shRNA construct together with GFP expression vectors. For the experimental control, we used neurons transfected only with GFP. The data are presented as mean ± SEM (n = 15 dendrites from 9 neurons for GFP control, n = 16 dendrites from 8 neurons for GFP plus control shRNA).

**Supplementary Figure 10.**
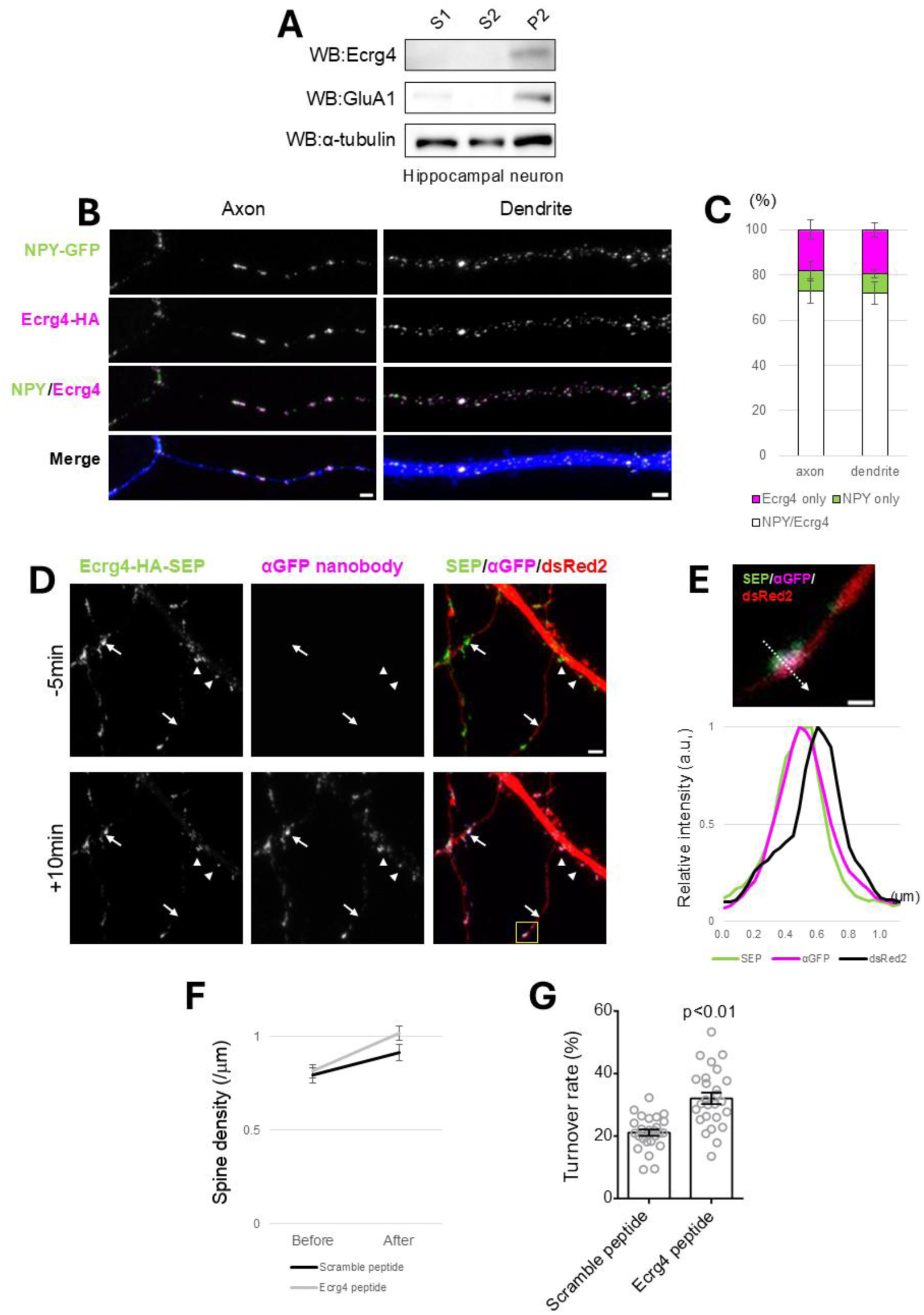
Expression and distribution of Ecrg4 in cultured hippocampal neurons. (A) Preferential localization of Ecrg4 protein in the membrane fraction. Immunoblotting of Ecrg4, GluA1AMPA-type glutamate receptor, and α-tubulin in the remaining fraction after removal of nuclei (S1), supernatant after centrifugation (S2), and resulting pellet (P2). (B) Images of an immunostained hippocampal neuron expressing HA-tagged Ecrg4 and NPY-GFP. The fluorescence signals were partially colocalized, suggesting Ecrg4 protein accumulation in dense-core vesicles. Bar = 2 μm. (C) Quantification of the overlap between puncta immunopositive for HA-tagged Ecrg4 and NPY-GFP. N = 4 cells from 1 dish. (D) Surface labeling with the anti-GFP nanobody revealing clusters of SEP-tagged Ecrg4 in dsRed2-positive axons (arrows) and dendrites (arrowheads). A nanobody was applied at time t = 0 min. Bar = 2 μm. (E) Enlarged image of a single Ecrg4 puncta in the axon, marked by a yellow square in (D), with the fluorescence intensity profile along the dashed arrow. Bar = 0.5 μm. (F) The spine density of cultured hippocampal neurons 24 hours after administration of either Ecrg4 peptides or scrambled peptides (n = 26 dendrites and 13 neurons for each condition). (G) Spine turnover rate of neurons treated with Ecrg4 peptides or scrambled peptides (n = 26 dendrites and 13 neurons for each condition).

## References

1. Rochefort, N.L., and Konnerth, A. (2012). Dendritic spines: from structure to in vivo function. EMBO Rep. 13, 699–708. 10.1038/EMBOR.2012.102.

2. Sala, C., and Segal, M. (2014). Dendritic spines: the locus of structural and functional plasticity. Physiol. Rev. 94, 141–188. 10.1152/physrev.00012.2013.

3. Matsuzaki, M., Ellis-Davies, G.C.R., Nemoto, T., Miyashita, Y., Iino, M., and Kasai, H. (2001). Dendritic spine geometry is critical for AMPA receptor expression in hippocampal CA1 pyramidal neurons. Nat. Neurosci. 4, 1086–1092. 10.1038/NN736.

4. Matsuzaki, M., Honkura, N., Ellis-Davies, G.C.R., and Kasai, H. (2004). Structural basis of long-term potentiation in single dendritic spines. Nature 429, 761–766. 10.1038/nature02617.

5. Nägerl, U.V., Eberhorn, N., Cambridge, S.B., and Bonhoeffer, T. (2004). Bidirectional activity-dependent morphological plasticity in hippocampal neurons. Neuron 44, 759–767. 10.1016/j.neuron.2004.11.016.

6. Zhou, Q., Homma, K.J., and Poo, M.M. (2004). Shrinkage of dendritic spines associated with long-term depression of hippocampal synapses. Neuron 44, 749–757. 10.1016/j.neuron.2004.11.011.

7. Loewenstein, Y., Kuras, A., and Rumpel, S. (2011). Multiplicative dynamics underlie the emergence of the log-normal distribution of spine sizes in the neocortex in vivo. Journal of Neuroscience 31, 9481–9488. 10.1523/JNEUROSCI.6130-10.2011.

8. Loewenstein, Y., Yanover, U., and Rumpel, S. (2015). Predicting the dynamics of network connectivity in the neocortex. Journal of Neuroscience 35, 12535–12544. 10.1523/JNEUROSCI.2917-14.2015.

9. Statman, A., Kaufman, M., Minerbi, A., Ziv, N.E., and Brenner, N. (2014). Synaptic size dynamics as an effectively stochastic process. PLoS Comput. Biol. 10. 10.1371/JOURNAL.PCBI.1003846.

10. Hazan, L., and Ziv, N.E. (2020). Activity dependent and independent determinants of synaptic size diversity. J. Neurosci. 40, 2828–2848. 10.1523/JNEUROSCI.2181-19.2020.

11. Tønnesen, J., Katona, G., Rózsa, B., and Nägerl, U.V. (2014). Spine neck plasticity regulates compartmentalization of synapses. Nat. Neurosci. 17, 678–685. 10.1038/nn.3682.

12. Kashiwagi, Y., Higashi, T., Obashi, K., Sato, Y., Komiyama, N.H., Grant, S.G.N., and Okabe, S. (2019). Computational geometry analysis of dendritic spines by structured illumination microscopy. Nat. Commun. 10, 1285. 10.1038/s41467-019-09337-0.

13. Zaccard, C.R., Shapiro, L., Martin-de-Saavedra, M.D., Pratt, C., Myczek, K., Song, A., Forrest, M.P., and Penzes, P. (2020). Rapid 3D enhanced resolution microscopy reveals diversity in dendritic spinule dynamics, regulation, and function. Neuron 107, 522–537. 10.1016/j.neuron.2020.04.025.

14. Penzes, P., Cahill, M.E., Jones, K.A., VanLeeuwen, J.-E., and Woolfrey, K.M. (2011). Dendritic spine pathology in neuropsychiatric disorders. Nat. Neurosci. 14, 285–293. 10.1038/nn.2741.

15. Fromer, M., Pocklington, A.J., Kavanagh, D.H., Williams, H.J., Dwyer, S., Gormley, P., Georgieva, L., Rees, E., Palta, P., Ruderfer, D.M., et al. (2014). De novo mutations in schizophrenia implicate synaptic networks. Nature 506, 179–184. 10.1038/NATURE12929.

16. Rodriguez-Murillo, L., Gogos, J.A., and Karayiorgou, M. (2012). The genetic architecture of schizophrenia: new mutations and emerging paradigms. Annu. Rev. Med. 63, 63–80. 10.1146/ANNUREV-MED-072010-091100.

17. Gulsuner, S., Walsh, T., Watts, A.C., Lee, M.K., Thornton, A.M., Casadei, S., Rippey, C., Shahin, H., Nimgaonkar, V.L., Go, R.C.P., et al. (2013). Spatial and temporal mapping of de novo mutations in schizophrenia to a fetal prefrontal cortical network. Cell 154, 518–529. 10.1016/J.CELL.2013.06.049.

18. Bourgeron, T. (2015). From the genetic architecture to synaptic plasticity in autism spectrum disorder. Nat. Rev. Neurosci. 16, 551–563. 10.1038/NRN3992.

19. Terashima, H., Minatohara, K., Maruoka, H., and Okabe, S. (2022). Imaging neural circuit pathology of autism spectrum disorders: autism-associated genes, animal models and the application of in vivo two-photon imaging. Microscopy (Oxf). 71, I81–I99. 10.1093/JMICRO/DFAB039.

20. Zieger, H.L., and Choquet, D. (2021). Nanoscale synapse organization and dysfunction in neurodevelopmental disorders. Neurobiol. Dis. 158, 105453. 10.1016/J.NBD.2021.105453.

21. Yilmaz, M., Yalcin, E., Presumey, J., Aw, E., Ma, M., Whelan, C.W., Stevens, B., McCarroll, S.A., and Carroll, M.C. (2021). Overexpression of schizophrenia susceptibility factor human complement C4A promotes excessive synaptic loss and behavioral changes in mice. Nat. Neurosci. 24, 214–224. 10.1038/S41593-020-00763-8.

22. Obi-Nagata, K., Suzuki, N., Miyake, R., MacDonald, M.L., Fish, K.N., Ozawa, K., Nagahama, K., Okimura, T., Tanaka, S., Kano, M., et al. (2023). Distorted neurocomputation by a small number of extra-large spines in psychiatric disorders. Sci. Adv. 9, eade5973. 10.1126/SCIADV.ADE5973.

23. Mukai, J., Dhilla, A., Drew, L.J., Stark, K.L., Cao, L., MacDermott, A.B., Karayiorgou, M., and Gogos, J.A. (2008). Palmitoylation-dependent neurodevelopmental deficits in a mouse model of 22q11 microdeletion. Nat. Neurosci. 11, 1302–1310. 10.1038/nn.2204.

24. Nakai, N., Takumi, T., Nakai, J., and Sato, M. (2018). Common defects of spine dynamics and circuit function in neurodevelopmental disorders: A systematic review of findings from in vivo optical imaging of mouse models. Front. Neurosci. 12. 10.3389/FNINS.2018.00412/PDF.

25. Shcheglovitov, A., Shcheglovitova, O., Yazawa, M., Portmann, T., Shu, R., Sebastiano, V., Krawisz, A., Froehlich, W., Bernstein, J.A., Hallmayer, J.F., et al. (2013). SHANK3 and IGF1 restore synaptic deficits in neurons from 22q13 deletion syndrome patients. Nature 503, 267–271. 10.1038/NATURE12618.

26. Llamosas, N., Arora, V., Vij, R., Kilinc, M., Bijoch, L., Rojas, C., Reich, A., Sridharan, B.P., Willems, E., Piper, D.R., et al. (2020). SYNGAP1 controls the maturation of dendrites, synaptic function, and network activity in developing human neurons. J. Neurosci. 40, 7980–7994. 10.1523/JNEUROSCI.1367-20.2020.

27. Yamamoto, K., Kuriu, T., Matsumura, K., Nagayasu, K., Tsurusaki, Y., Miyake, N., Yamamori, H., Yasuda, Y., Fujimoto, M., Fujiwara, M., et al. (2021). Multiple alterations in glutamatergic transmission and dopamine D2 receptor splicing in induced pluripotent stem cell-derived neurons from patients with familial schizophrenia. Transl. Psychiatry 11, 548. 10.1038/S41398-021-01676-1.

28. Wen, Z., Nguyen, H.N., Guo, Z., Lalli, M.A., Wang, X., Su, Y., Kim, N.S., Yoon, K.J., Shin, J., Zhang, C., et al. (2014). Synaptic dysregulation in a human iPS cell model of mental disorders. Nature 515, 414–418. 10.1038/NATURE13716.

29. Chisholm, K., Lin, A., Abu-Akel, A., and Wood, S.J. (2015). The association between autism and schizophrenia spectrum disorders: A review of eight alternate models of co-occurrence. Neurosci. Biobehav. Rev. 55, 173–183. 10.1016/J.NEUBIOREV.2015.04.012.

30. De Crescenzo, F., Postorino, V., Siracusano, M., Riccioni, A., Armando, M., Curatolo, P., and Mazzone, L. (2019). Autistic symptoms in schizophrenia spectrum disorders: A systematic review and meta-analysis. Front. Psychiatry 10. 10.3389/FPSYT.2019.00078/PDF.

31. Nenadić, I., Meller, T., Evermann, U., Schmitt, S., Pfarr, J.K., Abu-Akel, A., and Grezellschak, S. (2021). Subclinical schizotypal vs. autistic traits show overlapping and diametrically opposed facets in a non-clinical population. Schizophr. Res. 231, 32–41. 10.1016/J.SCHRES.2021.02.018.

32. Crespi, B., and Badcock, C. (2008). Psychosis and autism as diametrical disorders of the social brain. Behav. Brain Sci. 31, 241–261. 10.1017/S0140525X08004214.

33. Pratt, J., Winchester, C., Dawson, N., and Morris, B. (2012). Advancing schizophrenia drug discovery: optimizing rodent models to bridge the translational gap. Nat. Rev. Drug Discov. 11, 560–570. 10.1038/NRD3649.

34. Isshiki, M., Tanaka, S., Kuriu, T., Tabuchi, K., Takumi, T., and Okabe, S. (2014). Enhanced synapse remodelling as a common phenotype in mouse models of autism. Nat. Commun. 5, 4742. 10.1038/ncomms5742.

35. Lee, E., Lee, J., and Kim, E. (2017). Excitation/inhibition imbalance in animal models of autism spectrum disorders. Biol. Psychiatry 81, 838–847. 10.1016/J.BIOPSYCH.2016.05.011.

36. Tabuchi, K., Blundell, J., Etherton, M.R., Hammer, R.E., Liu, X., Powell, C.M., and Südhof, T.C. (2007). A neuroligin-3 mutation implicated in autism increases inhibitory synaptic transmission in mice. Science 318, 71–76. 10.1126/science.1146221.

37. Etherton, M., Földy, C., Sharma, M., Tabuchi, K., Liu, X., Shamloo, M., Malenka, R.C., and Südhof, T.C. (2011). Autism-linked neuroligin-3 R451C mutation differentially alters hippocampal and cortical synaptic function. Proc. Natl. Acad. Sci. U. S. A. 108, 13764–13769. 10.1073/pnas.1111093108.

38. Komiyama, N.H., Watabe, A.M., Carlisle, H.J., Porter, K., Charlesworth, P., Monti, J., Strathdee, D.J.C., O’Carroll, C.M., Martin, S.J., Morris, R.G.M., et al. (2002). SynGAP regulates ERK/MAPK signaling, synaptic plasticity, and learning in the complex with postsynaptic density 95 and NMDA receptor. The Journal of Neuroscience 22, 9721–9732. 22/22/9721 [pii].

39. Matsumura, K., Seiriki, K., Okada, S., Nagase, M., Ayabe, S., Yamada, I., Furuse, T., Shibuya, H., Yasuda, Y., Yamamori, H., et al. (2020). Pathogenic POGZ mutation causes impaired cortical development and reversible autism-like phenotypes. Nat. Commun. 11, 859. 10.1038/s41467-020-14697-z.

40. Nakatani, J., Tamada, K., Hatanaka, F., Ise, S., Ohta, H., Inoue, K., Tomonaga, S., Watanabe, Y., Chung, Y.J., Banerjee, R., et al. (2009). Abnormal behavior in a chromosome-engineered mouse model for human 15q11-13 duplication seen in autism. Cell 137, 1235–1246. 10.1016/j.cell.2009.04.024.

41. Baba, M., Yokoyama, K., Seiriki, K., Naka, Y., Matsumura, K., Kondo, M., Yamamoto, K., Hayashida, M., Kasai, A., Ago, Y., et al. (2019). Psychiatric-disorder-related behavioral phenotypes and cortical hyperactivity in a mouse model of 3q29 deletion syndrome. Neuropsychopharmacology 44, 2125–2135. 10.1038/s41386-019-0441-5.

42. Saito, R., Miyoshi, C., Koebis, M., Kushima, I., Nakao, K., Mori, D., Ozaki, N., Funato, H., Yanagisawa, M., and Aiba, A. (2021). Two novel mouse models mimicking minor deletions in 22q11.2 deletion syndrome revealed the contribution of each deleted region to psychiatric disorders. Mol. Brain 14. 10.1186/S13041-021-00778-7.

43. Nagahama, K., Sakoori, K., Watanabe, T., Kishi, Y., Kawaji, K., Koebis, M., Nakao, K., Gotoh, Y., Aiba, A., Uesaka, N., et al. (2020). Setd1a insufficiency in mice attenuates excitatory synaptic function and recapitulates schizophrenia-related behavioral abnormalities. Cell Rep. 32, 108126. 10.1016/J.CELREP.2020.108126.

44. Yamagata, Y., Kobayashi, S., Umeda, T., Inoue, A., Sakagami, H., Fukaya, M., Watanabe, M., Hatanaka, N., Totsuka, M., Yagi, T., et al. (2009). Kinase-dead knock-in mouse reveals an essential role of kinase activity of Ca^2+^/calmodulin-dependent protein kinase IIα in dendritic spine enlargement, long-term potentiation, and learning. Journal of Neuroscience 29, 7607–7618. 10.1523/JNEUROSCI.0707-09.2009.

45. Yamasaki, N., Maekawa, M., Kobayashi, K., Kajii, Y., Maeda, J., Soma, M., Takao, K., Tanda, K., Ohira, K., Toyama, K., et al. (2008). Alpha-CaMKII deficiency causes immature dentate gyrus, a novel candidate endophenotype of psychiatric disorders. Mol. Brain 1, 6. 10.1186/1756-6606-1-6.

46. Featherstone, R.E., Shimada, T., Crown, L.M., Melnychenko, O., Yi, J., Matsumoto, M., Tajinda, K., Mihara, T., Adachi, M., and Siegel, S.J. (2022). Calcium/calmodulin-dependent protein kinase IIα heterozygous knockout mice show electroencephalogram and behavioral changes characteristic of a subpopulation of schizophrenia and intellectual impairment. Neuroscience 499, 104–117. 10.1016/J.NEUROSCIENCE.2022.07.023.

47. Stephenson, J.R., Wang, X., Perfitt, T.L., Parrish, W.P., Shonesy, B.C., Marks, C.R., Mortlock, D.P., Nakagawa, T., Sutcliffe, J.S., and Colbran, R.J. (2017). A Novel Human CAMK2A Mutation Disrupts Dendritic Morphology and Synaptic Transmission, and Causes ASD-Related Behaviors. J. Neurosci. 37, 2216–2233. 10.1523/JNEUROSCI.2068-16.2017.

48. Brown, C.N., Cook, S.G., Allen, H.F., Crosby, K.C., Singh, T., Coultrap, S.J., and Bayer, K.U. (2021). Characterization of six CaMKIIα variants found in patients with schizophrenia. iScience 24. 10.1016/j.isci.2021.103184.

49. Qiu, S., Lu, Z., and Levitt, P. (2014). MET receptor tyrosine kinase controls dendritic complexity, spine morphogenesis, and glutamatergic synapse maturation in the hippocampus. J. Neurosci. 34, 16166–16179. 10.1523/JNEUROSCI.2580-14.2014.

50. Bloodgood, B.L., Sharma, N., Browne, H.A., Trepman, A.Z., and Greenberg, M.E. (2013). The activity-dependent transcription factor NPAS4 regulates domain-specific inhibition. Nature 503, 121–125. 10.1038/nature12743.

51. Spiegel, I., Mardinly, A.R., Gabel, H.W., Bazinet, J.E., Couch, C.H., Tzeng, C.P., Harmin, D.A., and Greenberg, M.E. (2014). Npas4 regulates excitatory-inhibitory balance within neural circuits through cell-type-specific gene programs. Cell 157, 1216–1229. 10.1016/J.CELL.2014.03.058.

52. Zamboni, V., Armentano, M., Saró, G., Ciraolo, E., Ghigo, A., Germena, G., Umbach, A., Valnegri, P., Passafaro, M., Carabelli, V., et al. (2016). Disruption of ArhGAP15 results in hyperactive Rac1, affects the architecture and function of hippocampal inhibitory neurons and causes cognitive deficits. Sci. Rep. 6. 10.1038/SREP34877.

53. Ba, W., Selten, M.M., van der Raadt, J., van Veen, H., Li, L.L., Benevento, M., Oudakker, A.R., Lasabuda, R.S.E., Letteboer, S.J., Roepman, R., et al. (2016). ARHGAP12 functions as a developmental brake on excitatory synapse function. Cell Rep. 14, 1355–1368. 10.1016/J.CELREP.2016.01.037.

54. Nakatani, Y., Kiyonari, H., and Kondo, T. (2019). Ecrg4 deficiency extends the replicative capacity of neural stem cells in a Foxg1-dependent manner. Development 146, dev168120. 10.1242/DEV.168120.

55. Richter, M., Lalli, E., and Ruggiero, C. (2023). Complex and pleiotropic signaling pathways regulated by the secreted protein augurin. Cell Commun. Signal. 21, 69. 10.1186/S12964-023-01090-8.

56. Chen, R., Liu, Y., Djekidel, M.N., Chen, W., Bhattacherjee, A., Chen, Z., Scolnick, E., and Zhang, Y. (2022). Cell type-specific mechanism of Setd1a heterozygosity in schizophrenia pathogenesis. Sci. Adv. 8, 1077. 10.1126/sciadv.abm1077.

57. Mukai, J., Cannavò, E., Crabtree, G.W., Sun, Z., Diamantopoulou, A., Thakur, P., Chang, C.Y., Cai, Y., Lomvardas, S., Takata, A., et al. (2019). Recapitulation and Reversal of Schizophrenia-Related Phenotypes in Setd1a-Deficient Mice. Neuron 104, 471–487.e12. 10.1016/j.neuron.2019.09.014.

58. MacDonald, M.L., Alhassan, J., Newman, J.T., Richard, M., Gu, H., Kelly, R.M., Sampson, A.R., Fish, K.N., Penzes, P., Wills, Z.P., et al. (2017). Selective loss of smaller spines in Schizophrenia. American Journal of Psychiatry 174, 586–594. 10.1176/APPI.AJP.2017.16070814/SUPPL_FILE/APPI.AJP.2017.16070814.DS001.PDF.

59. Pfeiffer, T., Poll, S., Bancelin, S., Angibaud, J., Inavalli, V.K., Keppler, K., Mittag, M., Fuhrmann, M., and Nägerl, U.V. (2018). Chronic 2P-STED imaging reveals high turnover of dendritic spines in the hippocampus in vivo. Elife 7. 10.7554/eLife.34700.

60. Attardo, A., Fitzgerald, J.E., and Schnitzer, M.J. (2015). Impermanence of dendritic spines in live adult CA1 hippocampus. Nature 523, 592–596. 10.1038/nature14467.

61. Moriguchi, T., Takeda, S., Iwashita, S., Enomoto, K., Sawamura, T., Koshimizu, U., and Kondo, T. (2018). Ecrg4 peptide is the ligand of multiple scavenger receptors. Sci. Rep. 8. 10.1038/S41598-018-22440-4.

62. Okabe, S., Kim, H.D., Miwa, A., Kuriu, T., and Okado, H. (1999). Continual remodeling of postsynaptic density and its regulation by synaptic activity. Nat. Neurosci. 2, 804–811. 10.1038/12175.

63. Shin, E., Kashiwagi, Y., Kuriu, T., Iwasaki, H., Tanaka, T., Koizumi, H., Gleeson, J.G., and Okabe, S. (2013). Doublecortin-like kinase enhances dendritic remodelling and negatively regulates synapse maturation. Nat. Commun. 4, 1440. 10.1038/ncomms2443.

64. Ooe, N., Saito, K., Mikami, N., Nakatuka, I., and Kaneko, H. (2004). Identification of a novel basic helix-loop-helix-PAS factor, NXF, reveals a Sim2 competitive, positive regulatory role in dendritic-cytoskeleton modulator drebrin gene expression. Mol. Cell. Biol. 24, 608–616. 10.1128/MCB.24.2.608-616.2004.

65. Ball, G., Demmerle, J., Kaufmann, R., Davis, I., Dobbie, I.M., and Schermelleh, L. (2015). SIMcheck: a toolbox for successful super-resolution structured illumination microscopy. Sci. Rep. 5, 15915. 10.1038/srep15915.

66. Love, M.I., Huber, W., and Anders, S. (2014). Moderated estimation of fold change and dispersion for RNA-seq data with DESeq2. Genome Biol. 15. 10.1186/S13059-014-0550-8.

